# Artificial intelligence aided design of peptides with custom secondary structure motifs and reduced amino acid alphabets

**DOI:** 10.64898/2026.04.29.721096

**Authors:** Sean M. Brown, Ashley B. Cohen, Scott N. Dean

## Abstract

Proteins are highly diverse functional polymers where the specific sequence of amino acids, selected from a standard genetically-encoded alphabet of twenty (C20), determines the structure and ultimately the function of the resulting folded protein. This standard alphabet has been identified to be non-randomly distributed in physicochemical properties crucial to both structure-formation and function, often referred to as coverage theory. While machine learning models have drastically improved protein structure prediction, success of protein design models lags structure prediction, particularly for custom secondary structure motifs and amino acid alphabets. Here we therefore bridge contemporary biological theory with recent advancements in artificial intelligence (AI) to develop and evaluate a generative AI protein design model, trained on hundreds of thousands of proteins within the RSCB PDB, for custom secondary structure motifs using reduced amino acid alphabets (RAAs). Results indicate an overall success in designing novel proteins with desired secondary structure motifs for a broad range of amino acid alphabets and complexity of designs. Interestingly, this tool often captures the full three-dimensional tertiary structure of a target protein despite training only on physicochemical sequence space and secondary structure information. The development of this model advances research across multiple disciplines, from general scientific AI architecture development to protein design for biotechnology, astrobiology, and early-Earth evolutionary biology.

## Introduction

All life on Earth since the last universal common ancestor (LUCA) has constructed metabolism primarily as a network of genetically encoded proteins.^1^ Proteins are highly diverse functional polymers composed of an amino acid sequence ‘selected’ from a standard 20-member genetically-encoded alphabet (C20). In 1972, Christian Anfinsen was awarded the Nobel Prize in Chemistry for demonstrating that a protein’s primary sequence (i.e., the specific order of amino acids) determines how a linear polymer folds into a three-dimensional conformation, ultimately defining a protein’s function.^2^

Since then, it has been identified that the alphabet of 20 genetically encoded amino acids exhibits a statistically non-random distribution, or *coverage,* in van der Waals’ volume (size), and hydrophobicity.^3–8^ Both size and hydrophobicity are critical to the formation of protein structure: hydrophobic collapse is widely accepted as an underlying principle,^9–11^ even the driver,^12^ of protein folding, while volume determines the steric constraints available to structure formation. Coverage is defined by the combination of the range of descriptors measured for a set of amino acids, and the evenness with which they are distributed across such range (see Fig. SI.2). While range allows for chemical diversity, evenness minimizes the phenotypic jump as one amino acid substitutes for another in any given sequence for any ‘desired’ three-dimensional structure. Recent work has even applied coverage theory to successfully design, synthesize, and characterize xeno amino acid sets with peptides bearing familiar secondary structures such as α-helices.^7^ We therefore sought to determine if these coverage-theory-grounded design principles could be leveraged to more readily design custom structural motifs using reduced amino acid alphabets (RAAs).

Due to recent advances in computational biology, research progress in canonical protein design has drastically accelerated over the last half decade. While *de novo* protein design has traditionally leveraged the reduction of sequence and feature space (e.g., Hecht binary patterning^13^), the use of artificial intelligence (e.g., ESM3^14^) to target a desired structure or function has emerged as the favored contemporary strategy. Recently developed models, largely with transformer architectures, are significantly more powerful than superannuated design methods.^15,16^ These models have proven capable of producing a wide variety of proteins with experimentally verified activity. For example, ProGen3 is a generative large language model that generates *de novo* protein sequences with similar properties to their naturally occurring analogs.^17^ Others, like ESM, can be utilized to simultaneously predict the structure and function of proteins via high quality embeddings.^14^ Additional critical developments here include protein generative diffusion models (e.g., RFdiffusion) for protein design^18^ and hybrid multimodal architectures that incorporate protein structure reinforcement learning to improve protein property predictions.^19^ These approaches enable the design of proteins with sequences and structures that are entirely novel compared to those found in nature. Recently, this expanded design space has been further extended by other newly developed computational tools, such as RareFold, which was developed for structure prediction and design of proteins containing xeno amino acids.^20^

As a contribution to this rapidly expanding protein design field, we developed and evaluated a generative artificial intelligence model (Fig. 1) to design novel proteins with custom secondary structure motifs and amino acid alphabets. Our model, which consists of a bidirectional Long Short-Term Memory architecture with multi-head self-attention (bLSTMa), was trained on a large, comprehensive dataset of secondary structure assignments (Dictionary of Secondary Structure in Proteins^21^, DSSP), associated physicochemical properties (van der Waals volume, logD, and formal charge at pH 7.0), density functional theory (DFT) derived potential energy surfaces (PES) for each amino acid, and amino acid backbone type (see Meringer et al., 2013^22^). Selection of volume, hydrophobicity, and charge is motivated primarily by the profound effect these two descriptors have on formation of protein secondary structure, alongside the apparent evolutionary optimization of these physicochemical descriptors.^3–5,8^ While many alternative molecular properties are available, our aim was to develop a computationally inexpensive, physics-grounded model. We explicitly avoid higher-dimensional property spaces such as homology, which may risk model overfitting and is antithetical to the *ab initio* nature of our method.

**Figure 1.**
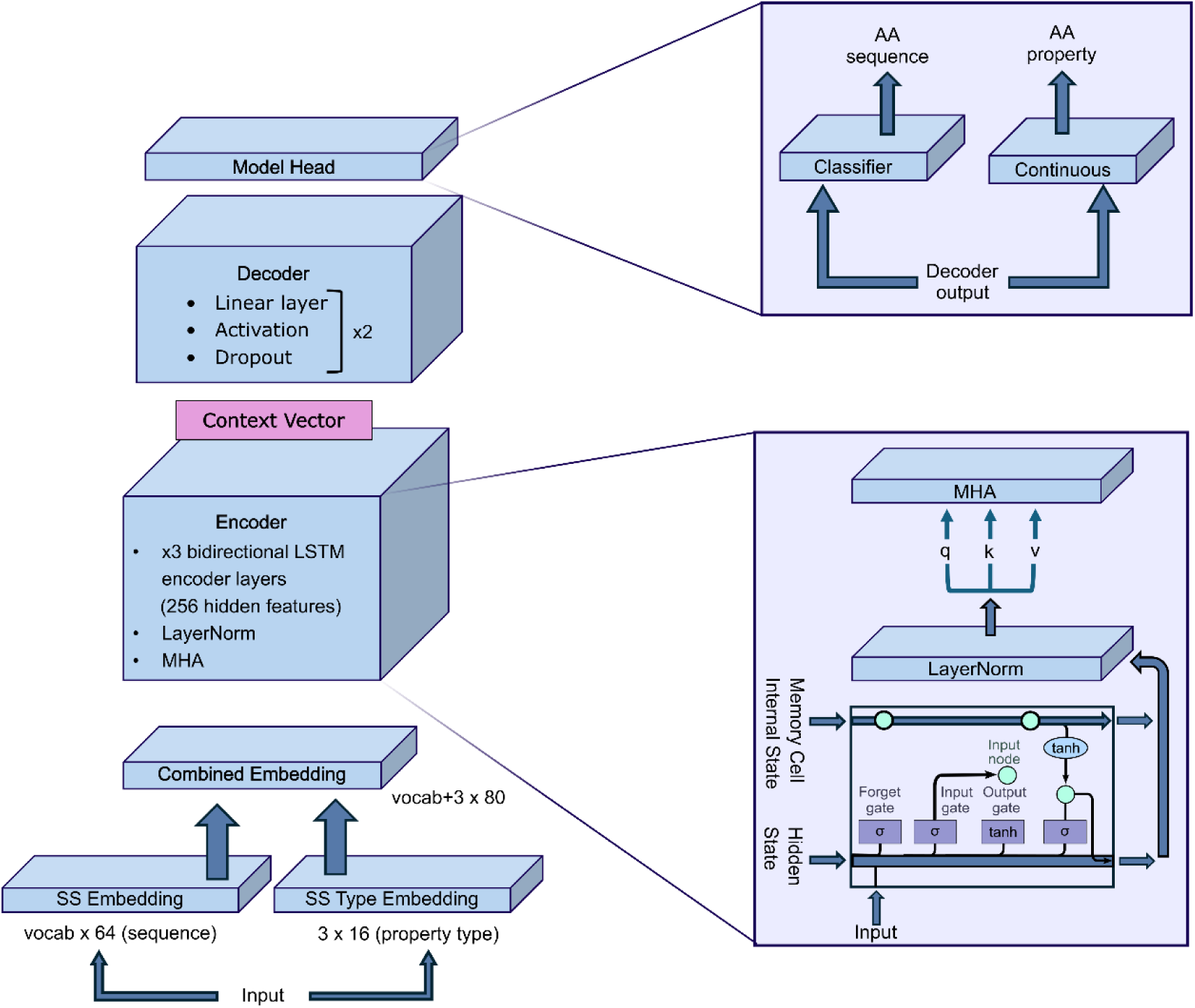
Major components of the bLSTMa encoder-decoder model architecture. Detailed architectures of the Encoder block, primarily made up of LSTM encoder layers and multi-head self-attention (bottom right) and model head, where the Decoder output is separately fed through a classifier and continuous value sequence to predict sequences and their associated properties (top right).

Utilizing the dataset of over two-hundred thousand DSSP-assigned protein structures within the RSCB PDB and the calculated properties at each residue, we designed and evaluated 62,600 sequences of proteins across 313 different amino acid alphabets to adopt numerous secondary structural motifs deriving from 200 target PDB structures. We evaluate the performance of our model by measuring accuracy metrics across various (i) protein lengths (ii) alphabet sizes (iii) structural diversity (iv) alphabet members and (v) three-dimensional structural propensity. Overall, results reveal that our model (Fig. 1) overall is successful in designing novel proteins with desired secondary structure motifs for a broad range of amino acid alphabets. Interestingly, many designs successfully retained the predicted targeted protein’s three-dimensional tertiary structure using RAAs, despite the lack of exposure to precise three-dimensional or atomistic-level information during training.

## Results

The alphabets used in this study derive from a comprehensive review of various reduced amino acid alphabets.^23^ This collection of alphabets comprises unique combinations of ∼83K alphabets resulting from 74 alphabet reduction methods. In order to reduce this large combinatorial alphabet space, we filtered this library to 312 alphabets using stratified sampling (see Table SI.1 and Methods).

To establish a baseline, we first measured background distributions of the amino acid alphabets used in this study. Figure 2A shows the distribution of unique RAAs by alphabet size, amino acid composition, and co-occurrence. We observe that the size of the alphabets identified in previous work^23^ follow a right-skewed normal distribution around a mean of 9.15 AAs and a standard deviation of 5.12 AAs. As expected, after filtering, the frequency of amino acids shifts to a uniform distribution (ranging from 6-19 AAs, x̅ = 11.94 AAs, and σ = 4.00 AAs). Furthermore, Figure 2B demonstrates that this uniformity did not come at the expense of altering the background amino acid frequency among these alphabets (which is also relatively uniform). We do observe a shift however in the filtered amino acid co-occurrence compared to the initial alphabet library (Fig. 2C).

**Figure 2.**
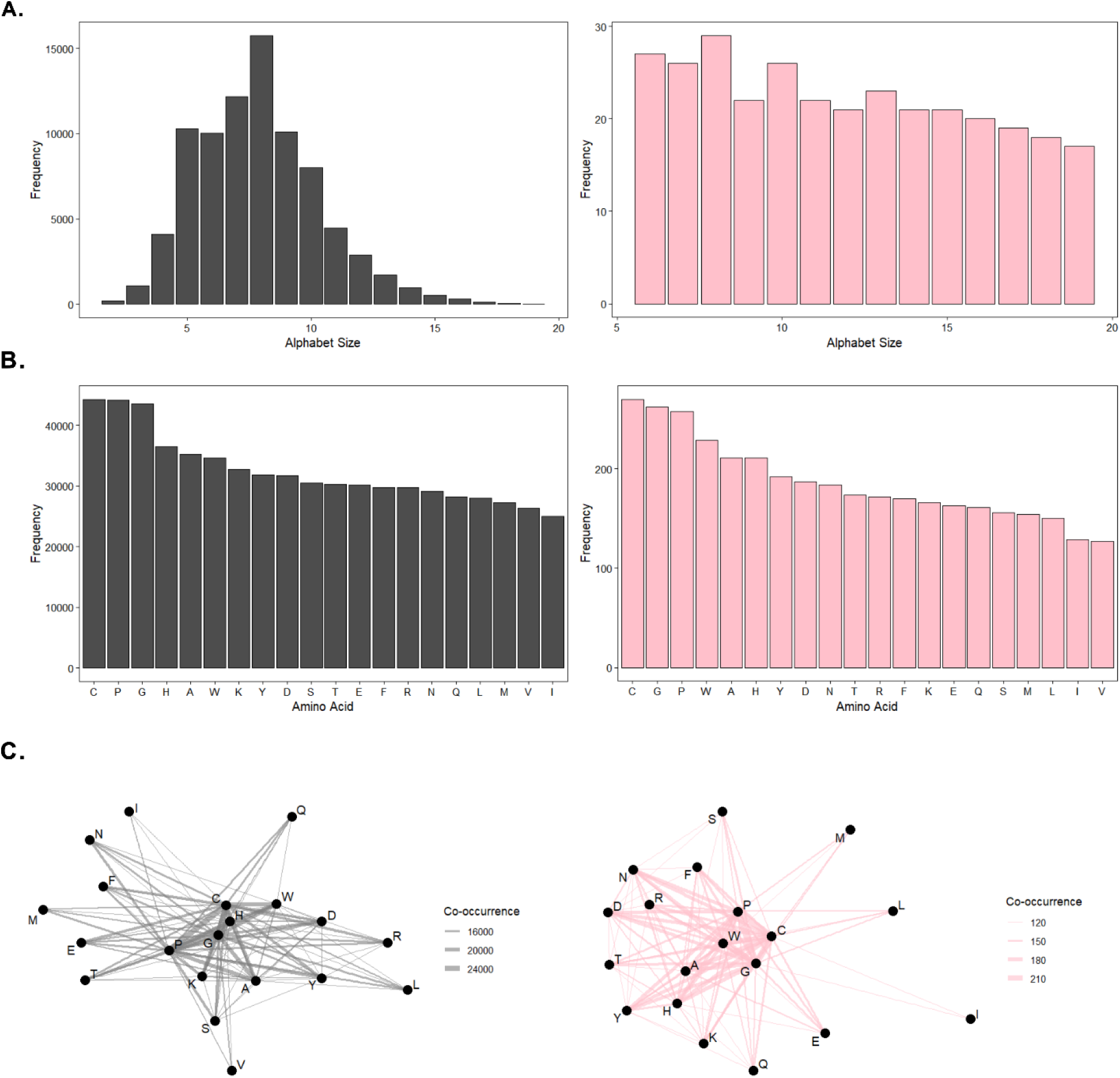
Background and subset alphabets characterization. **A.** distribution of background alphabets in terms of alphabet size for both the background ∼83k (grey) and subset ∼300 (pink) alphabets. **B.** distribution of background alphabets in terms of amino acid composition for both the background (grey) and subset (pink) alphabets. **C.** amino acid co-occurrence network for background alphabets for both the background (grey) and subset (pink) alphabets.

To develop a method that accurately generates amino acid sequences with desired secondary structure motifs, we trained a bLSTMa model on a diverse, non-redundant dataset of secondary structure assignments (DSSP) and associated physicochemical properties (van der Waals volume, logD, and formal charge at pH 7.0) extracted from >200K proteins within the RSCB PDB. From this set, by restricting chain length minimum and maximum sequence length to 40 and 250, respectively, we obtained a nonredundant set of 125,760 qualifying chains, from which randomly sampled 200 proteins as our evaluation set.

To evaluate the accuracy of the model in terms of predicted 3-state DSSP secondary structure (helix, sheet, coil), we calculated the three-state accuracy^24^ (Q3) of DSSP sequences to the target secondary structure (Fig. 3). After the model generates a sequence, we then predict secondary structure using ESMFold^25^, given the impracticality of synthesizing and characterizing thousands of generated sequences. For the preponderance of RAAs, the model displays a high degree of accuracy where deviation is primarily observed with RAAs fewer than 6 amino acids (Fig. 3A), where larger alphabets perform better in terms of Q3 compared to smaller alphabets (mean Q3 for size 19, 10, and 6 alphabets equals 87%, 72%, and 58%, respectively). We do however observe a sharp decline in performance (shifting from a left-skewed normal distribution to a right-skewed platykurtic distribution) for alphabets of size less than 10.

**Figure 3.**
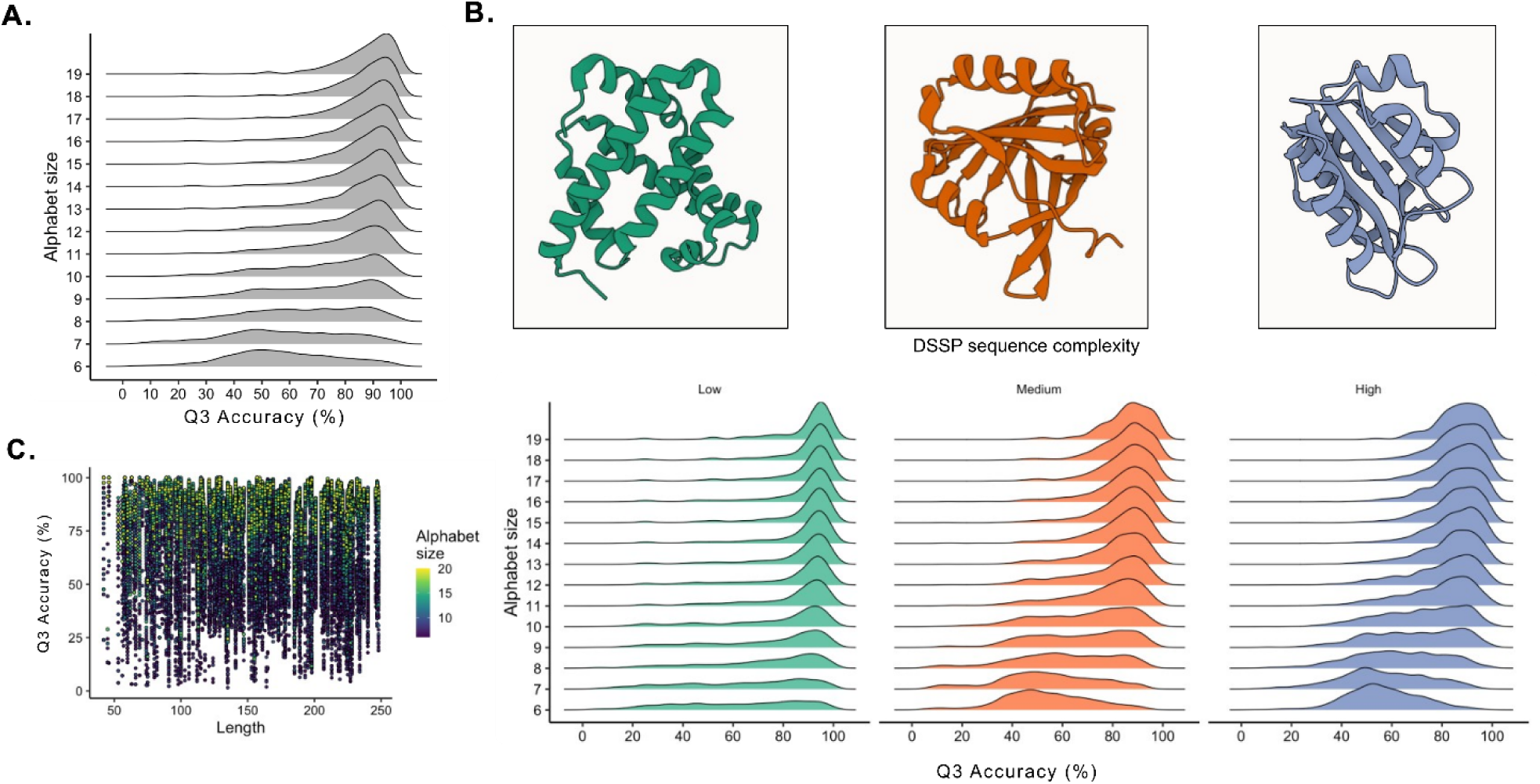
Model accuracy in terms of predicted 3-state DSSP secondary structure. **A.** Q3 accuracy of predicted sequences to target, input, secondary structure. **B.** Example target structures of low (green) medium (orange) and high (blue) structure complexity proteins (top) displayed in ribbon format. Histograms (bottom) of results for Q3 accuracy of predicted sequences to target, input, secondary structures for low- (green) medium- (orange) and high- (blue) complexity protein targets. **C.** Q3 accuracy for predicted sequences across protein sequence length and alphabet size.

We suspect the rapid decline in performance for alphabets comprising fewer than ten amino acids is due to a general inability to capture sufficient physicochemical diversity within the available chemical space. With increasingly fewer amino acids comes fewer degrees of freedom in each potential sequence, thereby increasing the difficulty of generating a sequence with the desired motif. This is especially true with increasingly more complex design motifs, which we observe less frequently captured when constructing sequences with alphabets comprising few amino acids. Lack of diversity within chemical space precluding complex secondary structure additionally aligns well with early-Earth evolutionary literature, which suggests primordial, reduced, amino acid alphabets likely functionally and structurally-relied on cofactors (e.g. metal ions) for structure and catalysis.^26–29^ Although the average Q3 decreased with alphabet size, interestingly the maximum did not. In other words, while predominantly less abundant, accurate DSSP designs are indeed present for every alphabet tested; while for most alphabets tested (size 19-10) the model demonstrated strong performance in terms of Q3.

The Q3 accuracy of the generated sequences does appear to partially depend on the overall complexity of the target structure (Fig. 3B). Low-complexity structures, such as only α-helix, are therefore more abundant when using RAAs compared to higher complexity structures (e.g., a helix-coil-sheet-coil-sheet-coil-helix motif). As structural complexity increases, we generally observe a shift from a left-tailed distribution (low complexity, mean Q3 = 79%) to a slightly platykurtic distribution (medium complexity, mean Q3 = 73%; high complexity, mean Q3 = 75%). The effect of alphabet size is observed here as well. For instance, across all design complexities, alphabets sized 19-10 vastly outperform smaller alphabets (Fig. 3B). While far less common than designs with low-complexity structures, we do however achieve high Q3 designs with high-complexity structure using alphabets as small as 6 amino acids. Furthermore, we tested model accuracy as a function of protein sequence length. We do not observe dependence on sequence length for model accuracy (measured by target DSSP Q3, Fig. 3C; R^2^ = -0.03), therefore demonstrating that our method works equally across a broad size-range of target structures (Fig. 3C).

Beyond the overall predictive accuracy of target secondary structure, we additionally investigated if our model inadvertently learned underlying principles which govern the three-dimensional folding of amino acid chains. To do so, we measured the extent to which three-dimensional (tertiary) structure of target proteins is preserved in our model’s output design constructs. Remarkably, despite having only been trained on the one-dimensional information of secondary structure sequence motifs and associated physicochemical properties, the model’s output often appears to capture precise tertiary structure of the PDB target (Fig. 4). As expected, this outcome appears to depend upon which amino acid alphabet is provided to our model. Specifically, we observe a clear relationship between amino acid alphabet size and three-dimensional structural accuracy where an increase in alphabet size typically improves both TM- and pLDDT-scores while also lowering the backbone RMSD (Fig. 4A). There is a noticeable shift in TM- and pLDDT-scores from a bimodal to a low-accuracy normal unimodal distribution between 8-11 amino acid alphabets. For RMSD, we observe a right-skewed platykurtic shift to a left-skewed distribution as the alphabet size decreases. This finding suggests heavyweight models and immense datasets may not always be required, depending on the target sequence and design complexity because of the relationship shown between particular secondary structure design constructs and predicted tertiary structure. Furthermore, the consistency with observed DSSP Q3 accuracy indicates that training merely on secondary structures and sequence-dependent physicochemical data appears to be sufficient for accurate three-dimensional design rather than relying on extracted patterns within life’s amino acid sequence-structure space (i.e., homology modeling – as in traditional models).

**Figure 4.**
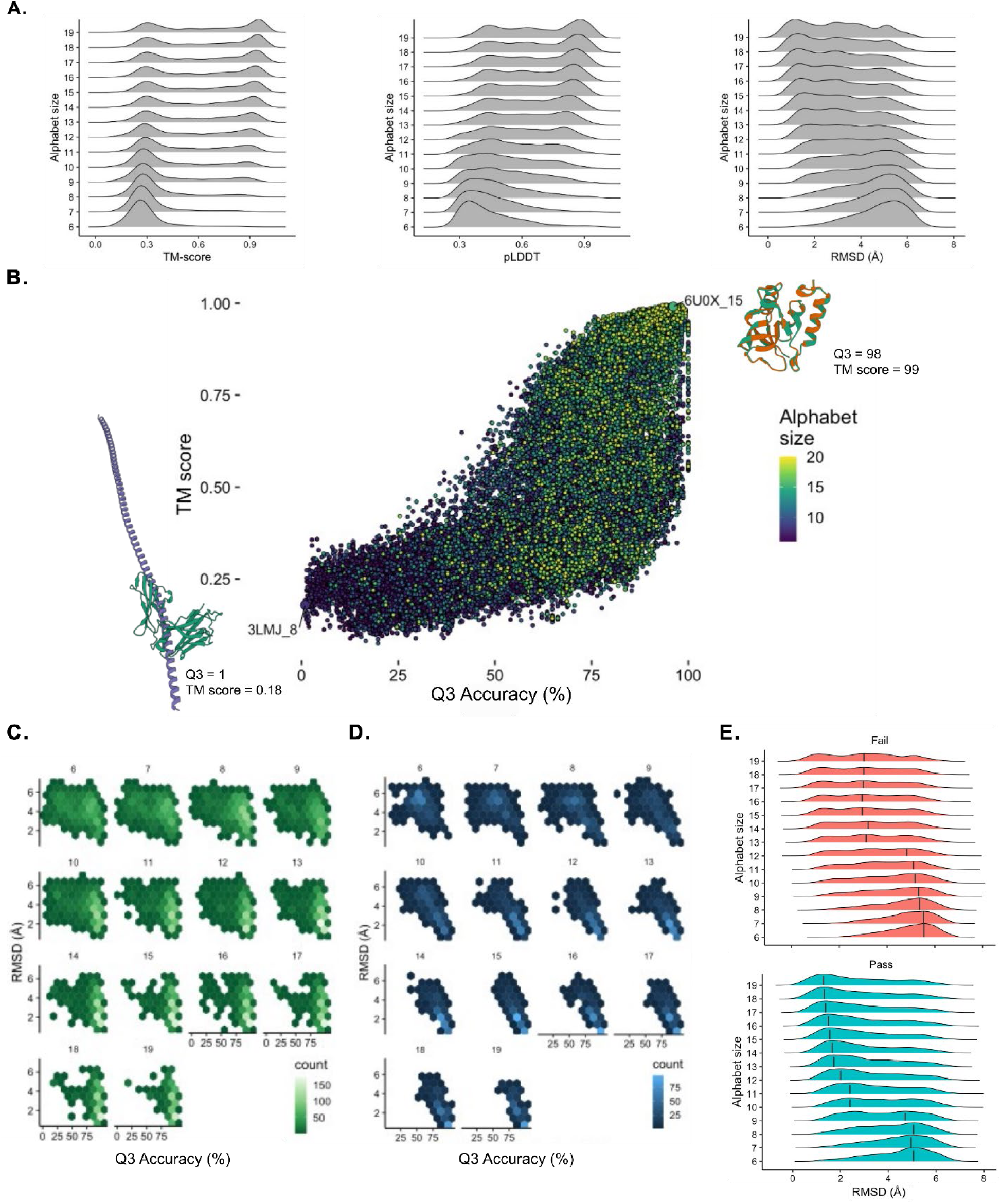
Model accuracy in terms of target tertiary structure. **A.** Accuracy for various amino acid alphabet sizes (6-19) measured by ESMFold TM-score (left), pLDDT (middle), and RMSD (right). **B. Q3** and pLDDT scores across various alphabet sizes (6-19). Lowest (3LMJ_8) and highest (6U0X_15) accuracy to target structures (green – target; orange – ESMFold predicted model output sequence) measured by pLDDT and Q3 shown in bottom left and top right respectively. **C-D.** RMSD versus Q3 accuracy correlation for low (**C.** - green) and high (**D. -** blue) structure complexity sequences. **E.** RMSD of designs stratified by alphabet size and either Pass or Fail after being passed through the designability model. Density curves are shown with modal outcomes highlighted with a black bar.

To further investigate this relationship and the features governing design success, we analyzed the relationship between DSSP sequence properties and Q3. A random forest model trained on DSSP and sequence-derived features was able to accurately predict whether a design would have high (>80%) or low (≤ 80%) Q3 score, achieving a receiver operating characteristic area under the curve (ROC-AUC) of 0.81 (Fig. SI.1A). This indicates that design accuracy is strongly determined by the properties of the target secondary structure sequence. Feature-importance analysis of this model revealed that properties such as proportion of helix and Simpson diversity were highly predictive of accuracy (Fig. SI.1B), again, suggesting that complex DSSP sequences are less likely to yield high-Q3 scoring designs. These features were then extracted into a designability model (see Methods). We passed each DSSP in the training set through this model and categorized into either a ‘pass’ or ‘fail’ using a prediction probability threshold of 0.6. The resulting RMSDs of the passing and failed show a significant separation at certain alphabet sizes (Fig. 4E). Specifically, the modal difference in RMSD for alphabets of size ≥ 10 was > 1 Å, with the greatest difference observed for alphabets of size 10-12 each with a difference of ∼2.8 Å, and the modal passing design with alphabet size ≥ 13 had a RMSD of < 2 Å (Fig. 4E), demonstrating that this simple random forest model can be used as pre-filter to significantly improve downstream design outcomes.

While these metrics (TM, pLDDT, and RMSD) of three-dimensional accuracy generally concur, we do not observe as strong of a relationship for measures of three-dimensional accuracy, Q3, and alphabet size (Fig. 4B). Furthermore, we do not observe a strong correlation between Q3 and pLDDT (R^2^ = 0.65). While the more accurate predictions were those using larger alphabets and the poor accuracy predictions were typically observed with smaller alphabets, the space between these two extremes is far more complex. For example, the most accurate construct observed (6U0X_15; pLDDT = 0.98; Q = 99%) derives from a 15-mer alphabet; therefore, underscoring how high-accuracy constructs (in both DSSP and three-dimensional structure as predicted by ESMFold) are available beyond the very largest alphabets. Moreover, Figure 4C-D shows that high-accuracy constructs (particularly for larger RAAs) also retained a relatively low RMSD to the three-dimensional structure of the original target in PDB. While this pattern is true for high-complexity sequences (Fig. 4D) it is, perhaps unsurprisingly, stronger for low complexity sequences (Fig. 4C). The probability of this phenomenon occurring for a more complex protein however would intuitively be lower than with a less complex structure (e.g., only α-helix). One plausible explanation of this finding is that there are physical constraints on particular secondary structure patterns which result in only one possible three-dimensional conformation at physiological conditions. Alternatively, an equally likely explanation may be artifacts deriving from the training data for ESMFold structure prediction; where the only available data of course was all of life’s proteins and therefore ESMFold could not have plausibly learned other ways to form certain three-dimensional patterns. At present, the sheer lack of experimentally resolved constructs built with alphabets beyond C20 (N=1, Xeno Peptide P2^7^) obscures where the truth lies between these two explanations.

## Discussion

In this work we developed a bLSTMa machine learning model, trained on a large, diverse, non-redundant dataset of secondary structure assignments (DSSP) and associated physicochemical properties. This model can be used as a physicochemically-grounded protein design tool for many desired secondary structural motifs and nearly any canonical amino acid alphabet. Moreover, since the training data derives from physicochemistry rather than biological presuppositions (such as homology), it is plausible that this model can be extended for design of proteins comprising entirely synthetic, xeno, amino acid alphabets.

To benchmark our model for canonical RAAs, we therefore designed and evaluated many proteins comprising various canonical reduced amino acid alphabets to adopt the same secondary structural motifs observed in 200 target PDB protein structures. Results demonstrate that our model is preponderantly able to design proteins with custom secondary structure motifs with many RAAs based solely on fundamental physicochemical properties of amino acids. When coupled with a designability model, we further improve accessibility at identification of successful design motifs using RAAs as few as 6 AAs reliably. The development of this tool advances research progress across multiple disciplines, including general scientific AI architecture development, protein design for biotechnology, astrobiology, and early-Earth evolutionary biology.

### Implications for artificial intelligence and machine learning

As the field of biology-focused artificial intelligence continues to rapidly grow, vision, language, and multimodal AI models have become larger and more complex, often having hundreds of millions to tens of billions of trainable parameters.^30^ While larger models often exhibit superior performance, in practice, they are expensive to develop and difficult to deploy given the enormous computational resources required (ibid.). By contrast, to sufficiently train our 5.1 million parameter model, merely 70% of data entries need to be seen. Even with small batching (16 entries), training can be completed in ≤ 40 epochs and 18 hours wall time on a Nvidia A100 GPU, overall achieving a relatively low computational training cost. Furthermore, we demonstrate that our model successfully generates protein sequences with numerous amino acid alphabets at state-of-the-art predicted performance using a simple LSTM encoder-decoder architecture with a modest number of trainable parameters and small vocabulary size. Our computationally accessible architecture therefore allows for greater user accessibility, more precise architecture refinement and scaling, faster training, and richer contextual embeddings.

As such, we are sanguine that a broader user-developer community will iteratively refine and improve upon this method (e.g., dead node pruning, adjusting the context window size, changing the number of encoder layers, etc.) and tune for each particular use-case. Furthermore, the bLSTMa is trained on inherently more enriched embeddings compared to a standard LLM. Most LLMs feed a single unidirectional embedding through their architecture as a consequence of being generative transformers (i.e., autoregressive, single-direction, prediction of new tokens from prior tokens). The model used here instead concatenates bidirectional embeddings of secondary structure and physicochemical properties into a highly enriched combined contextual embedding. The low-loss and reduced training overhead of bidirectional embeddings has been demonstrated by the popularity and success of a similar approach: ProteinBERT.^31^ While quick and easy to use, ProteinBERT only predicts protein properties, while the model demonstrated here is for generative protein design of particular secondary structure motifs.

### Implications for protein design

Our approach advances protein design by establishing a first approximation of a minimum number of physicochemical descriptors needed to reliably generate protein sequences based solely on a desired sequence of secondary structural motifs. Rather than exhaustively including a large dataset of numerous physicochemical descriptors, we built our framework on one set of “essential” parameters through optimality and evolutionary amino acid alphabet coverage theories. This simple, physicochemically grounded approach keeps the model’s vocabulary size small, which directly determines the size of the model weights’ last tensor dimension. A small tensor size not only decreases training time, but also drastically reduces the overhead memory requirements to train the model. This is particularly interesting when compared to protein language models like ProGen2, which pre-train using all available taxonomy, function, and ontogeny ids associated with proteins to create prepended conditional “tags”, resulting in an enormous vocabulary.

The novelty of this approach is not limited to just life’s amino acids. Very few tools exist for xeno peptide and protein research apart from recently developed approaches like RareFold^32^ given the lack of data for entirely noncanonical peptides/proteins^7^ when compared to the vast repertoire provided by life’s amino acid alphabet. Because our bLSTMa was trained using position-dependent physicochemical properties and secondary structure information instead of individual amino acids themselves, our model is, in theory, plausibly extensible to any amino acid alphabet, including entirely xeno amino acid alphabets. The accuracy of structures predicted from xeno amino acid alphabets is of course yet to be determined due to the fundamental lack of xeno peptides or protein experimental data to date. In other words, it is uninformative to assess the performance of a polymer-class where the sample size is miniscule. The approach presented here therefore guides any synthetic biologist to efficiently design proteins or peptides to a high degree of confidence with any canonical amino acid alphabet while not excluding compatibility for xeno amino acid polymers. Furthermore, as more xeno peptides and proteins are synthetically developed, those structures can be used to test, and subsequently fine-tune this model for xeno amino acid reliability. As this framework is adapted and built upon, it will become clearer how many and which physicochemical parameters are needed to most accurately distill the physics of protein secondary structure and to what extent additional physicochemistry leads to diminishing returns regarding training and prediction. Until then, our model may usefully serve as a proof-of-concept starting point for a small-vocabulary, physicochemically based secondary structure design tool.

### Implications for astrobiology and evolutionary biology

The method presented in this study additionally advances the fields of astrobiology and evolutionary biology. Our bLSTMa acts as a novel framework for scientists to design and test theorized primordial or xeno protein candidates. With the model presented here, and specifically with the ability to confidently design structures from RAAs, we present a new foundation with which to probe evolutionary history of proteins prior to LUCA. Similarly, this approach informs how readily an alternative origin of life (bearing protein biochemistry) may emerge by enabling the beginnings to answer: could alternative amino acid alphabets form eerily similar proteins with entirely different building blocks?

## Conclusions

Our model trained on a limited subset of physicochemical parameters and our benchmarking protocol demonstrates that a spartan physics and chemistry-based approach can indeed be used to design and generate proteins with various amino acid alphabets and desired secondary structures. Many other physicochemical parameters exist that may or may not improve the model’s ability to design peptides or proteins. The physicochemical combinatorial space is simply too vast to be completely explored. It is therefore a tractable near-future research direction to begin traversing that space by testing the extent to which other physicochemical properties contribute to the formation of fundamental protein secondary structure. Additionally, it remains unknown if and to what extent a similar small-parameter model could successfully generate structurally and *functionally* novel protein and peptide sequences. While this model eases the design of custom secondary structures using RAAs, substanital progress here ultimately hinges on future work conducting experimental synthesis, characterization, and validation of novel synthetic constructs.

## Methods

### Library Curation

Our machine learning model was trained on all proteins within the RCSB PDB (www.rcsb.org) protein-only group (as of 01JAN2026, N=236,183). For each protein, the DSSP^21,33^ sequence was extracted if available, or calculated, and a sequence of physicochemical descriptors (van der Waals volume, logD, and formal charge at pH 7.0; Table 1), together with three per-residue Ramachandran basin free energies (α-helix, β-sheet, and polyproline-II), were assigned to each residue in each sequence, determined by using the potential energy surfaces described below. Molecular volume and hydrophobicity were calculated using ChemAxon JChem, https://www.chemaxon.com, (see Mayer-Bacon and Yirk^34^ for an in-depth explanation on how to calculate these descriptors).

**Table 1.** Physicochemical value for each genetically encoded amino acid at pH 7.0.

| Amino Acid<br>(1-Letter) | Van Der Waals<br>Volume ($\text{\AA}^3$) | JChem<br>LogD | Formal<br>Charge |
| --- | --- | --- | --- |
| A | 86.4 | -1.34 | 0 |
| C | 104.9 | -1.29 | 0 |
| D | 118.6 | -1.98 | -1 |
| E | 135.9 | 1.70 | -1 |
| F | 159.0 | 0.31 | 0 |
| G | 69.1 | -1.91 | 0 |
| H | 131.7 | -2.11 | 0 |
| I | 138.3 | -0.01 | 0 |
| K | 149.3 | -1.48 | +1 |
| L | 138.3 | -0.09 | 0 |
| M | 139.5 | -0.69 | 0 |
| N | 120.8 | -2.79 | 0 |
| P | 108.6 | -1.07 | 0 |
| Q | 138.1 | -2.50 | 0 |
| R | 168.6 | -2.54 | +1 |
| S | 95.2 | -2.39 | 0 |
| T | 112.5 | -1.97 | 0 |
| V | 121 | -0.46 | 0 |
| W | 178.7 | 0.41 | 0 |
| Y | 167.8 | 0.01 | 0 |

### Potential Energy Surfaces

We calculated the potential energy surface (PES) for each genetically encoded amino acid with acetylated N-termini and N-methylamidated C-termini as previously described in detail^7,35–37^. Capping the backbone termini avoid artificial electrostatics and charge effects while simultaneously more closely emulating the behavior of each AA as when in a protein. Calculations were conducted with consideration of each amino acid in its native protonation state in water at pH 7.0.

In order to generate potential energy surfaces, we first generated initial conformers for every pair (N=144) of phi and psi dihedral angles (-180, -150, -120, -90, -60, -30, 0, 30, 60, 90, 120, 150), for each AA, with random starting 3D geometries generated in RDKit (v.2024.03.5, https://www.rdkit.org) before optimization with the UFF general force field^38^ (iteratively increasing force constraints: initial = 0.002, multiplier = 2.0 per iteration) for every desired phi and psi dihedral angles within a five-degree tolerance. UFF-optimized conformers were further optimized in xTB at a GFN2-xTB^39^ with ɸ, Ψ dihedral angle constraints (force constant = 0.05) in ALPB implicit solvation.^40^ The Conformer–Rotamer Ensemble Sampling Tool^41^ (CREST) program was then used to thoroughly sample each amino acid side chain for each conformation, while constraining phi and psi angles to respective target values (force constant = 0.25). Finally, DFT single point energies were calculated in Orca^42^ (v.6.1.0) for every CREST conformer returned within 10 kcal⋅mol ^-1^, with openCOSMO-RS,^43^ BP86 functional,^44^ def2-TZVP basis set,^45^ and Grimme’s D3(BJ) dispersion correlations.^46^ Final free energies of conformers (G) were calculated with the formula:

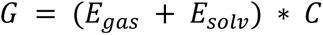

Where the final free energy (*G*) is equal to the sum of free energy of gas phase (*E_gas_*) and solvation (*E_solv_*) multiplied by the Hartree-to-kcal⋅mol^-1^ constant (*C*). The structure with the lowest DFT(BP86-D3BJ)/def2-TZVP//openCOSMO-RS energy was considered as the representative conformer for that phi and psi angle combination. The lowest DFT(BP86-D3BJ)/def2-TZVP//openCOSMO-RS energy across the entire PES for each amino acid was considered the relative minima for that amino acid and all other energies across the PES were scaled appropriately to that value. The PES data for these twenty amino acids and previous PES calculations can be found in prior work^7,35–37^.

### Design Space Selection

We downselected from the 82,795 distinct reduced canonical amino acid alphabets in Liang et al.^23^ to 312 by performing stratified sampling after removing alphabets of size 5 or smaller, grouping by alphabet size and method (as defined by Liang et al.) and randomly selecting one per group (Table SI.1). To obtain evaluation targets, we randomly sampled 200 protein chains from the curated PDB dataset, restricting to chains of 40-250 residues (125,760 qualifying chains; median length = 155 residues). The full design space consisted of 200 targets by 313 alphabets (312 reduced + 1 full) = 62,600 predicted sequences, all of which were folded with ESMFold and scored.

### Machine Learning Model and Training

The dataset was randomly split 70/20/10 into training, validation, and test sets, batched into groups of 16. DSSP sequences were tokenized into a vocabulary of nine states (α-helix, 3_10_-helix, π-helix, β-strand, bridge, turn, bend, polyproline, coil) and embedded as 64-dimensional vectors, concatenated with a 16-dimensional secondary structure type embedding (helix/sheet/coil), yielding an 80-dimensional combined input.

Our model has an encoder-decoder architecture featuring bidirectional LSTM layers and a multi-head self-attention mechanism (Fig. 1), totaling 5,065,280 trainable parameters. The combined embedding first passes through the encoder block, which consists of three bidirectional long short-term memory (LSTM) encoder layers with 256 hidden features, layer normalization and eight-head multi-head attention. The resulting context vector is then passed into the decoder block, which consists of two iterations of a linear layer, rectified linear unit activation function and dropout sequence. The model was trained for up to 40 epochs with early stopping (patience = 5) using AdamW (learning rate = 0.001, weight decay = 0.01) with a cosine annealing warm restart schedule (T₀ = 20, T_mult = 2). Temperature was linearly annealed from 2.0 to 0.5 over training to transition from exploratory to exploitative optimization. The loss function combined three terms: property MSE (α = 1.0) between the predicted and target six-dimensional per-residue descriptor (van der Waals volume, logD, and formal charge at pH 7.0, plus α-helix, β-sheet, and polyproline-II Ramachandran basin free energies), cross-entropy with teacher forcing (β = 0.2), and entropy regularization (γ = 0.01), with an additional L2 penalty on logits. Gradient clipping (max norm = 1.0) and mixed precision training were used for stability and efficiency. The classifier output layer weights were initialized with Xavier Uniform (gain = 0.1) and zero bias.

During inference, the model outputs logits over all 20 amino acids at each position. For reduced alphabet predictions, logits corresponding to disallowed amino acids are set to negative infinity before applying softmax, effectively restricting the output distribution to only the permitted alphabet. The amino acid with the highest probability at each position is then selected (greedy decoding).

### Structure Prediction and Validation

Predicted protein sequences were folded using ESMFold (facebook/esmfold_v1), run locally as a single-sequence predictor (no MSA), with PyTorch 2.6 (CUDA 12.4), transformers 5.12, and Python 3.11. The predicted local distance difference test (pLDDT) was extracted from ESMFold output for each prediction as a measure of fold confidence. Secondary structure of each predicted structure was assigned with DSSP (mkdssp v4.0). To assess structural accuracy, predicted structures were compared to experimentally resolved target structures from the RCSB PDB using TM-align, yielding RMSD and TM-score metrics.

### Performance Quantification

To evaluate the range of target sequence complexity over which the design method is effective, we computed a composite complexity score for each DSSP sequence from three metrics: (1) Shannon entropy, measuring the diversity of secondary structure states in the sequence; (2) normalized transition frequency, measuring the rate of secondary structure type changes along the sequence normalized by maximum possible transitions; and (3) Simpson diversity index, measuring the evenness of secondary structure state usage. We rescaled each metric to fall between 0 and 1, where Shannon entropy was divided by log_2_(*N*) for the maximum entropy of the DSSP states, and the composite complexity score was computed as the average of the three metrics. Target sequences were grouped into three equally sized complexity categories (low, medium, and high) using quantile-based binning to examine the relationship between target complexity and design accuracy.

So as to quantify model accuracy, we measured the three-state accuracy (Q3) of the predicted DSSP sequence relative to the target DSSP sequence across alphabet sizes and target complexities. As a metric for three-dimensional accuracy, we additionally measured the RMSD of predicted output structures to the experimentally resolved target structures found in the RCSB PDB.

### Designability Classifier

To identify which target structures are amenable to reduced-alphabet design, we trained a classifier to distinguish high- from low-designability targets, defined as either > 80% or ≤ 80% Q3 score. Input features were derived from each target’s DSSP: the fraction of residues in each DSSP state (9-state, and the collapsed 3-state helix/strand/coil), the longest contiguous helix and strand segments, and three secondary-structure complexity metrics (normalized Shannon entropy, transition frequency, and Simpson diversity) and their composite. Several model families (L1/L2/elastic-net logistic regression, linear SVM, random forest, and gradient boosting) were compared by repeated stratified five-fold cross-validation optimizing ROCAUC. A minimal model using only the three-state helix/strand/coil fractions and their composite complexity achieved comparable performance (ROCAUC 0.79). Native-sequence-derived features (amino-acid composition and physicochemical descriptors computed with Biopython) were additionally evaluated. Analyses were implemented in Python with scikit-learn (v1.8).

## Acknowledgements

S.M.B. and A.B.C. acknowledge a postdoctoral fellowship through the National Research Council’s Research Associateship Program. This research was funded by the Office of Naval Research (WU# 61A1P4). The opinions and assertions contained herein are those of the authors and are not to be construed as those of the U.S. Navy, military service at large or the U.S. Government. S.N.D. is an employee of the U.S. Government.

## Author Contributions

Conceptualization: S.M.B., A.B.C., S.N.D..; Computational investigation: S.M.B., A.B.C., S.N.D.; Formal analysis: S.M.B., S.N.D.; Data curation: S.M.B., S.N.D.; Visualization: S.M.B., A.B.C., S.N.D.; Writing– original draft: S.M.B., A.B.C., S.N.D.; Writing–editing: S.M.B., A.B.C., S.N.D.; Supervision: S.N.D.; Project administration: S.M.B., S.N.D.; Funding acquisition: S.N.D.

## Data Availability Statement

All data needed to evaluate the conclusions in the paper are present in the paper, the Supplementary Information. Any additional data is available by request.

