## Supplementary Information for "Artificial intelligence aided design of peptides with custom secondary structure motifs and reduced amino acid alphabets"

FOR

DISTRIBUTION STATEMENT A. Approved for public release: distribution is unlimited.

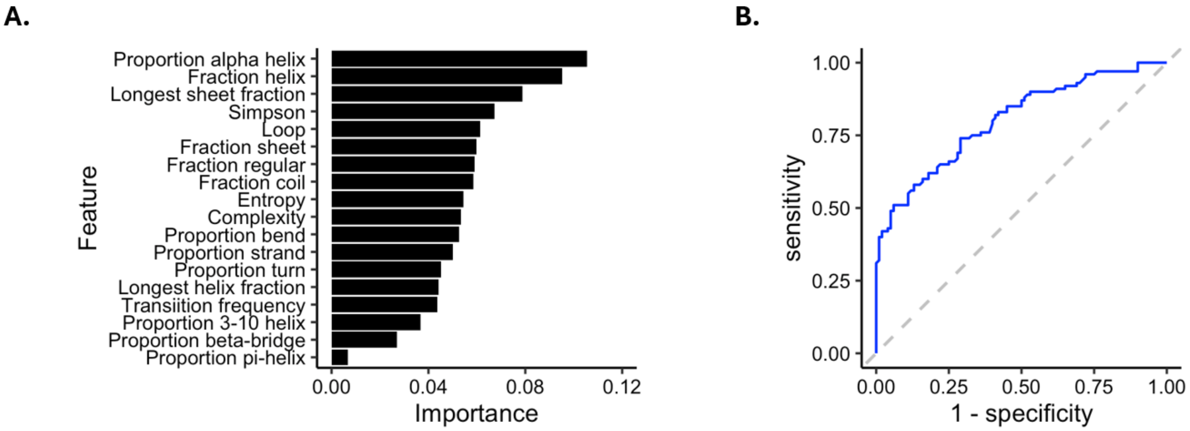


**Figure SI.1. DSSP percent identity prediction using sequence-based features. A**. Receiver operating characteristic curve (ROC) of the best-performing classifier. ROC-AUC = 0.98. **B.** Top 18 features ranked by importance for the Random Forest classifier.

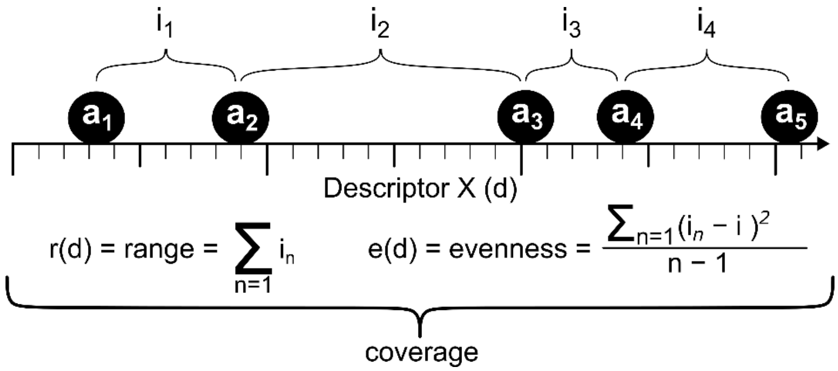


**Figure SI.2.** Physicochemical coverage - definition of range and evenness. For a given chemical descriptor such as van der Waals volume, "coverage" joins two statistics to characterize a set of amino acids. These statistics are shown for an example set of five amino acids ($a_1 \dots a_5$) with four corresponding intervals ($i_1 \dots i_4$) measured in terms of the hypothetical quantitative 'descriptor x' ($d$). Evenness ($e$) is the sample variance of the intervals between amino acids ($i$); Range ($r$) is the sum of these intervals ($\sum i_{1 \dots 4}$); "Coverage" is therefore the combination of range AND evenness for any given physicochemical descriptor.

**Table SI.1 Stratified Reduced Amino Acid Alphabets**

| Method | Size | Reduced Code | Combination |
|---|---|---|---|
| AIS | 6 | SWGA-DRHQYNE-KLVIF-C-PM-T | W-D-L-C-M-T |
| AIS | 7 | SQLGE-DRP-KT-W-HN-C-VIMAFY | G-R-K-W-H-C-Y |
| AIS | 8 | DKP-SRGFNE-WH-C-Q-LVMAY-I-T | P-S-W-C-Q-A-I-T |
| AIS | 9 | SGA-DP-RWHYN-KE-C-Q-LT-IV-MF | A-P-N-E-C-Q-L-V-M |
| BLOSUM40 | 5 | AGPST-RNDQEHK-C-ILMFYV-W | G-D-C-M-W |
| BLOSUM40 | 6 | AGPST-RNDQEK-C-H-ILMFYV-W | A-E-C-H-M-W |
| BLOSUM40 | 7 | ANDGST-RQEK-C-H-ILMFYV-P-W | G-Q-C-H-L-P-W |
| BLOSUM40 | 8 | ANDGST-RQEK-C-H-ILMV-FY-P-W | G-Q-C-H-M-Y-P-W |
| BLOSUM40 | 9 | AGST-RQEK-ND-C-H-ILMV-FY-P-W | T-Q-D-C-H-I-F-P-W |
| BLOSUM40 | 10 | AGST-RK-ND-C-QE-H-ILMV-FY-P-W | S-R-D-C-E-H-L-Y-P-W |
| BLOSUM40 | 11 | AST-RK-ND-C-QE-G-H-ILMV-FY-P-W | A-R-N-C-E-G-H-V-F-P-W |
| BLOSUM40 | 12 | AST-RK-ND-C-QE-G-H-IV-LM-FY-P-W | S-K-N-C-Q-G-H-V-L-Y-P-W |
| BLOSUM40 | 13 | AST-RK-N-D-C-QE-G-H-IV-LM-FY-P-W | A-K-N-D-C-Q-G-H-V-M-F-P-W |
| BLOSUM40 | 14 | AST-RK-N-D-C-Q-E-G-H-IV-LM-FY-P-W | S-R-N-D-C-Q-E-G-H-I-M-F-P-W |
| BLOSUM40 | 15 | A-RK-N-D-C-Q-E-G-H-IV-LM-FY-P-ST-W | A-R-N-D-C-Q-E-G-H-V-L-F-P-T-W |
| BLOSUM40 | 16 | A-RK-N-D-C-Q-E-G-H-IV-LM-F-P-ST-W-Y | A-K-N-D-C-Q-E-G-H-V-L-F-P-S-W-Y |
| BLOSUM40 | 17 | A-R-N-D-C-Q-E-G-H-IV-LM-K-F-P-ST-W-Y | A-R-N-D-C-Q-E-G-H-V-L-K-F-P-T-W-Y |
| BLOSUM40 | 18 | A-R-N-D-C-Q-E-G-H-IV-LM-K-F-P-S-T-W-Y | A-R-N-D-C-Q-E-G-H-I-M-K-F-P-S-T-W-Y |
| BLOSUM40 | 19 | A-R-N-D-C-Q-E-G-H-IV-L-K-M-F-P-S-T-W-Y | A-R-N-D-C-Q-E-G-H-I-L-K-M-F-P-S-T-W-Y |
| BLOSUM50 | 5 | IMVL-FWY-G-PCAST-NHQEDRK | V-W-G-P-H |
| BLOSUM50 | 6 | LVIM-AGST-PHC-FYW-EDNQ-KR | V-S-C-Y-N-K |
| BLOSUM50 | 7 | IMVL-FWY-G-P-CAST-NHQED-RK | V-Y-G-P-A-Q-R |
| BLOSUM50 | 8 | LVIMC-AG-ST-P-FYW-EDNQ-KR-H | M-G-S-P-F-E-K-H |
| BLOSUM50 | 9 | IMV-L-FWY-G-P-C-AST-NHQED-RK | M-L-F-G-P-C-A-D-K |
| BLOSUM50 | 10 | IMV-L-FWY-G-P-C-A-STNH-QERK-D | I-L-F-G-P-C-A-T-E-D |
| BLOSUM50 | 11 | IMV-L-FWY-G-P-C-A-STNH-QRK-E-D | V-L-F-G-P-C-A-S-R-E-D |
| BLOSUM50 | 12 | LVIM-C-A-G-ST-P-FY-W-EQ-DN-KR-H | I-C-A-G-T-P-Y-W-E-N-K-H |
| BLOSUM50 | 13 | IMV-L-F-WY-G-P-C-A-ST-N-HQRK-E-D | M-L-F-W-G-P-C-A-S-N-K-E-D |
| BLOSUM50 | 14 | IMV-L-F-WY-G-P-C-A-S-T-N-HQRK-E-D | V-L-F-Y-G-P-C-A-S-T-N-K-E-D |
| BLOSUM50 | 15 | IMV-L-F-WY-G-P-C-A-S-T-N-H-QRK-E-D | M-L-F-W-G-P-C-A-S-T-N-H-K-E-D |
| BLOSUM50 | 16 | IMV-L-F-W-Y-G-P-C-A-S-T-N-H-QRK-E-D | V-L-F-W-Y-G-P-C-A-S-T-N-H-R-E-D |
| BLOSUM50 | 18 | LM-VI-C-A-G-S-T-P-F-Y-W-E-D-N-Q-K-R-H | M-I-C-A-G-S-T-P-F-Y-W-E-D-N-Q-K-R-H |
| BLOSUM62 | 5 | FWYH-MILV-CATSP-G-NQDERK | Y-L-A-G-K |
| BLOSUM62 | 6 | FWYH-MILV-CATS-P-G-NQDERK | Y-I-A-P-G-Q |
| BLOSUM62 | 7 | FWYH-MILV-CATS-P-G-NQDE-RK | W-M-T-P-G-D-R |
| BLOSUM62 | 8 | FWYH-MILV-CA-NTS-P-G-DE-QRK | W-I-C-S-P-G-D-K |
| BLOSUM62 | 9 | FWYH-ML-IV-CA-NTS-P-G-DE-QRK | H-L-I-C-S-P-G-E-K |
| BLOSUM62 | 10 | FWY-ML-IV-CA-TS-NH-P-G-DE-QRK | W-L-V-C-T-N-P-G-E-R |
| BLOSUM62 | 11 | FWY-ML-IV-CA-TS-NH-P-G-D-QE-RK | W-M-I-C-T-H-P-G-D-E-R |
| BLOSUM62 | 12 | FWY-ML-IV-C-A-TS-NH-P-G-D-QE-RK | Y-M-V-C-A-T-N-P-G-D-E-R |

| | | | |
|---|---|---|---|
| BLOSUM62 | 13 | FWY-ML-IV-C-A-T-S-NH-P-G-D-QE-RK | F-L-V-C-A-T-S-H-P-G-D-Q-R |
| BLOSUM62 | 15 | FWY-ML-IV-C-A-T-S-N-H-P-G-D-QE-R-K | Y-L-I-C-A-T-S-N-H-P-G-D-Q-R-K |
| BLOSUM62 | 16 | W-FY-ML-IV-C-A-T-S-N-H-P-G-D-QE-R-K | W-Y-M-I-C-A-T-S-N-H-P-G-D-E-R-K |
| BLOSUM62 | 17 | W-FY-ML-IV-C-A-T-S-N-H-P-G-D-Q-E-R-K | W-Y-L-I-C-A-T-S-N-H-P-G-D-Q-E-R-K |
| BLOSUM62 | 18 | W-FY-M-L-IV-C-A-T-S-N-H-P-G-D-Q-E-R-K | W-F-M-L-V-C-A-T-S-N-H-P-G-D-Q-E-R-K |
| BLOSUM62 | 19 | W-F-Y-M-L-IV-C-A-T-S-N-H-P-G-D-Q-E-R-K | W-F-Y-M-L-V-C-A-T-S-N-H-P-G-D-Q-E-R-K |
| BLOSUM62 & MC | 5 | CFYW-MLIV-G-PATS-NHQEDRK | C-L-G-A-K |
| BLOSUM62 & MC | 6 | CFYW-MLIV-G-P-ATS-NHQEDRK | Y-V-G-P-T-D |
| BLOSUM62 & MC | 7 | CFYW-MLIV-G-P-ATS-NHQED-RK | C-V-G-P-A-E-K |
| BLOSUM62 & MC | 8 | CFYW-MLIV-G-P-ATS-NH-QED-RK | Y-V-G-P-S-N-E-R |
| BLOSUM62 & MC | 9 | CFYW-ML-IV-G-P-ATS-NH-QED-RK | F-M-V-G-P-S-H-E-R |
| BLOSUM62 & MC | 10 | C-FYW-ML-IV-G-P-ATS-NH-QED-RK | C-W-M-I-G-P-T-H-E-R |
| BLOSUM62 & MC | 11 | C-FYW-ML-IV-G-P-A-TS-NH-QED-RK | C-Y-L-V-G-P-A-T-N-E-R |
| BLOSUM62 & MC | 12 | C-FYW-ML-IV-G-P-A-TS-NH-QE-D-RK | C-F-M-V-G-P-A-S-N-Q-D-K |
| BLOSUM62 & MC | 13 | C-FYW-ML-IV-G-P-A-T-S-NH-QE-D-RK | C-Y-L-V-G-P-A-T-S-H-Q-D-K |
| BLOSUM62 & MC | 14 | C-FYW-ML-IV-G-P-A-T-S-N-H-QE-D-RK | C-F-M-V-G-P-A-T-S-N-H-Q-D-K |
| BLOSUM62 & MC | 15 | C-FYW-ML-IV-G-P-A-T-S-N-H-QE-D-R-K | C-W-L-V-G-P-A-T-S-N-H-Q-D-R-K |
| BLOSUM62 & MC | 16 | C-FY-W-ML-IV-G-P-A-T-S-N-H-QE-D-R-K | C-Y-W-M-I-G-P-A-T-S-N-H-E-D-R-K |
| BLOSUM62 & MC | 17 | C-FY-W-ML-IV-G-P-A-T-S-N-H-Q-E-D-R-K | C-F-W-M-I-G-P-A-T-S-N-H-Q-E-D-R-K |
| BLOSUM62 & MC | 18 | C-FY-W-M-L-IV-G-P-A-T-S-N-H-Q-E-D-R-K | C-Y-W-M-L-V-G-P-A-T-S-N-H-Q-E-D-R-K |
| BLOSUM62 & MC | 19 | C-F-Y-W-M-L-IV-G-P-A-T-S-N-H-Q-E-D-R-K | C-F-Y-W-M-L-V-G-P-A-T-S-N-H-Q-E-D-R-K |
| Boltzmann relation | 5 | VILMFWYA-C-ED-RK-GPSTHQN | M-C-E-R-G |
| Boltzmann relation | 6 | VILMFWY-A-C-ED-RK-GPSTHQN | I-A-C-E-R-H |
| Boltzmann relation | 7 | VILMFWY-A-C-ED-RK-G-PSTHQN | W-A-C-D-K-G-P |
| Boltzmann relation | 8 | VILMF-WY-A-C-ED-RK-G-PSTHQN | L-W-A-C-E-K-G-S |
| Boltzmann relation | 9 | VILMF-WY-A-C-ED-RK-G-P-STHQN | F-W-A-C-D-R-G-P-H |
| Boltzmann relation | 10 | VILMF-WY-A-C-ED-RK-G-P-ST-HQN | L-Y-A-C-D-K-G-P-T-H |
| Boltzmann relation | 14 | VILM-F-W-Y-A-C-ED-R-K-G-P-ST-H-QN | V-F-W-Y-A-C-E-R-K-G-P-S-H-N |
| Chemistry properties | 8 | DE-KRH-QN-ST-P-CM-WYF-GALIV | E-R-N-S-P-C-Y-A |
| Chemistry properties | 10 | DE-N-KRH-Q-T-SGAC-P-YWF-M-LIV | E-N-R-Q-T-G-P-Y-M-V |
| Chemistry space | 5 | NQHRK-YFW-MC-STDEGAPVL-I | N-W-M-L-I |
| Clustering analysis | 5 | APST-CGHIL-DENQ-FMWY-KRV | A-G-E-F-V |
| Clustering analysis | 6 | APST-CILMV-DENQ-FWY-G-HKR | S-L-Q-W-G-K |
| Contact potential | 5 | DE-KR-NQSTGPHY-ACW-FMLIV | D-R-G-C-F |
| Contact potential | 7 | DENQ-KR-G-P-AV-STHWY-CFMLI | N-K-G-P-V-Y-L |
| Distance matrix | 5 | LMWFY-CIV-NPHST-AG-DEQRK | M-V-N-G-D |
| Distance matrix | 7 | ND-HST-GPC-WFY-IVLM-RK-AQE | N-S-G-W-L-K-Q |
| Distance matrix | 8 | LM-WFY-CIV-NP-HST-AG-DE-QRK | L-Y-I-N-S-A-E-K |
| Distance matrix | 12 | LM-W-FY-C-IV-NP-H-ST-AG-DE-Q-RK | L-W-Y-C-V-P-H-T-A-E-Q-R |
| Dynamic Programming | 10 | DE-KRH-NQ-ST-ILV-FWY-C-M-AG-P | E-H-Q-S-L-Y-C-M-A-P |
| Dynamic clustering | 5 | TVMLFW-C-YA-G-RNDQEHKPST | M-C-A-G-T |
| Dynamic clustering | 6 | TVMLF-WY-C-AH-G-RNDQEKPST | M-Y-C-H-G-Q |
| Dynamic clustering | 7 | TVMLF-WY-C-AH-GP-R-NDQEKST | M-W-C-H-P-R-K |

| | | | |
|---|---|---|---|
| Dynamic clustering | 8 | TVMLF-WY-C-A-G-Q-R-NHDEKPST | F-W-C-A-G-Q-R-T |
| Dynamic clustering | 9 | TVMLF-WY-C-A-G-P-H-K-RQNDEST | L-W-C-A-G-P-H-K-N |
| Dynamic clustering | 10 | TVML-F-W-Y-C-A-H-G-RN-QPKDEST | V-F-W-Y-C-A-H-G-R-S |
| Dynamic clustering | 11 | TVMLF-W-Y-C-A-H-G-R-N-Q-PKDEST | L-W-Y-C-A-H-G-R-N-Q-S |
| Dynamic clustering | 12 | TVML-F-W-Y-C-A-H-G-N-Q-T-RDEKPS | L-F-W-Y-C-A-H-G-N-Q-T-S |
| Dynamic clustering | 13 | TVML-F-W-Y-C-A-H-G-R-N-Q-P-DEKST | L-F-W-Y-C-A-H-G-R-N-Q-P-D |
| Dynamic clustering | 14 | TVML-F-W-Y-C-A-H-G-R-N-Q-P-K-DEST | V-F-W-Y-C-A-H-G-R-N-Q-P-K-T |
| Dynamic clustering | 15 | TVML-F-W-Y-C-A-H-G-R-N-Q-P-K-D-EST | T-F-W-Y-C-A-H-G-R-N-Q-P-K-D-T |
| Dynamic clustering | 16 | TVML-F-W-Y-C-A-H-G-R-N-Q-P-K-S-T-DE | T-F-W-Y-C-A-H-G-R-N-Q-P-K-S-T-D |
| Dynamic clustering | 17 | TVL-M-F-W-Y-C-A-H-G-R-N-Q-P-K-S-T-DE | V-M-F-W-Y-C-A-H-G-R-N-Q-P-K-S-T-D |
| Dynamic clustering | 18 | TVL-M-F-W-Y-C-A-H-G-R-N-Q-P-K-S-T-D-E | T-M-F-W-Y-C-A-H-G-R-N-Q-P-K-S-T-D-E |
| ECGA | 5 | ACIY-MFLV-GHTN-SWDE-PRKQ | I-M-T-W-P |
| Fuzzy clustering | 5 | EHKQDNILPTACGMS-V-FY-R-W | Q-V-Y-R-W |
| Fuzzy clustering | 6 | DEHIKLMNPQRST-FWY-A-C-G-V | M-F-A-C-G-V |
| Fuzzy clustering | 7 | ADEGHIKLNPQST-C-M-R-V-FY-W | H-C-M-R-V-Y-W |
| Fuzzy clustering | 8 | ADEGNPQST-HIKL-R-V-FY-C-M-W | S-K-R-V-F-C-M-W |
| Fuzzy clustering | 10 | DEHIKLNPQT-FY-M-R-S-W-A-C-G-V | L-Y-M-R-S-W-A-C-G-V |
| Fuzzy clustering | 11 | DENQT-HIKLR-FY-M-P-S-W-A-C-G-V | Q-K-F-M-P-S-W-A-C-G-V |
| Fuzzy clustering | 13 | EHKQ-DN-IL-PT-FY-M-R-S-W-A-C-G-V | Q-D-I-T-Y-M-R-S-W-A-C-G-V |
| Fuzzy clustering | 14 | EHKQ-DN-IL-PT-A-C-G-M-S-V-F-Y-R-W | H-N-L-T-A-C-G-M-S-V-F-Y-R-W |
| Fuzzy clustering | 15 | EQ-D-N-T-HIKL-R-FY-M-P-S-W-A-C-G-V | Q-D-N-T-L-R-Y-M-P-S-W-A-C-G-V |
| Fuzzy clustering | 16 | EQ-H-K-DN-IL-P-T-FY-M-R-S-W-A-C-G-V | Q-H-K-N-I-P-T-F-M-R-S-W-A-C-G-V |
| Fuzzy clustering | 17 | EQ-H-K-DN-IL-P-T-M-R-S-F-Y-W-A-C-G-V | Q-H-K-D-L-P-T-M-R-S-F-Y-W-A-C-G-V |
| Fuzzy clustering | 18 | E-Q-D-N-T-IL-H-K-R-FY-M-P-S-W-A-C-G-V | E-Q-D-N-T-I-H-K-R-Y-M-P-S-W-A-C-G-V |
| Fuzzy clustering | 19 | E-H-K-Q-D-N-IL-P-T-A-C-G-M-S-V-F-Y-R-W | E-H-K-Q-D-N-L-P-T-A-C-G-M-S-V-F-Y-R-W |
| GDM | 5 | AG-CDQ-EHILNVYFMRSW-K-PT | G-D-H-K-P |
| GDM | 7 | AG-CDQ-EHNY-FMRSW-ILV-K-PT | A-D-N-S-V-K-P |
| GDM | 8 | AG-C-DQ-EHNY-FMRSW-ILV-K-PT | A-C-D-E-M-V-K-P |
| GDM | 9 | AG-C-DQ-EHNY-FMW-ILV-K-PT-RS | A-C-D-Y-W-V-K-T-R |
| GDM | 10 | A-C-DQ-EHNY-FMW-G-ILV-K-PT-RS | A-C-D-H-F-G-I-K-P-R |
| GDM | 11 | A-C-DQ-EHNY-FM-G-ILV-K-PT-RS-W | A-C-Q-H-M-G-V-K-P-S-W |
| GDM | 12 | A-C-DQ-EHNY-FM-G-IL-K-PT-RS-V-W | A-C-Q-N-F-G-L-K-P-R-V-W |
| GDM | 13 | A-C-DQ-E-FM-G-HNY-IL-K-PT-RS-V-W | A-C-Q-E-M-G-Y-I-K-P-S-V-W |
| GDM | 14 | A-C-D-E-FM-G-HNY-IL-K-PT-Q-RS-V-W | A-C-D-E-M-G-H-I-K-T-Q-S-V-W |
| GDM | 15 | A-C-D-E-FM-G-HNY-IL-K-PT-Q-R-S-V-W | A-C-D-E-M-G-N-I-K-T-Q-R-S-V-W |
| GDM | 16 | A-C-D-E-F-G-HNY-IL-K-M-PT-Q-R-S-V-W | A-C-D-E-F-G-Y-L-K-M-P-Q-R-S-V-W |
| GDM | 17 | A-C-D-E-F-G-HNY-IL-K-M-P-Q-R-S-T-V-W | A-C-D-E-F-G-Y-L-K-M-P-Q-R-S-T-V-W |
| GDM | 18 | A-C-D-E-F-G-HNY-I-K-L-M-P-Q-R-S-T-V-W | A-C-D-E-F-G-H-I-K-L-M-P-Q-R-S-T-V-W |
| GDM | 19 | A-C-D-E-F-G-HN-I-K-L-M-P-Q-R-S-T-V-W-Y | A-C-D-E-F-G-N-I-K-L-M-P-Q-R-S-T-V-W-Y |
| GONNET | 5 | W-C-GPHNDERQKAST-FY-VMIL | W-C-A-F-L |
| GONNET | 6 | W-C-G-PHNDERQKAST-FY-VMIL | W-C-G-E-F-I |
| GONNET | 7 | W-C-G-P-HNDERQKAST-FY-VMIL | W-C-G-P-D-F-L |
| GONNET | 8 | W-C-G-P-H-NDERQKAST-FY-VMIL | W-C-G-P-H-N-Y-L |

| | | | |
|---|---|---|---|
| GONNET | 9 | W-C-G-P-H-NDERQK-AST-FY-VMIL | W-C-G-P-H-N-T-F-L |
| GONNET | 10 | W-C-G-P-H-NDERQK-AST-F-Y-VMIL | W-C-G-P-H-E-A-F-Y-L |
| GONNET | 11 | W-C-G-P-H-NDE-RQK-AST-F-Y-VMIL | W-C-G-P-H-D-Q-S-F-Y-I |
| GONNET | 12 | W-C-G-P-H-N-DE-RQK-AST-F-Y-VMIL | W-C-G-P-H-N-E-Q-S-F-Y-L |
| GONNET | 13 | W-C-G-P-H-N-DE-RQK-AST-F-Y-V-MIL | W-C-G-P-H-N-E-K-A-F-Y-V-M |
| GONNET | 14 | W-C-G-P-H-N-D-E-RQK-AST-F-Y-V-MIL | W-C-G-P-H-N-D-E-K-A-F-Y-V-L |
| GONNET | 15 | W-C-G-P-H-N-D-E-R-QK-AST-F-Y-V-MIL | W-C-G-P-H-N-D-E-R-K-T-F-Y-V-L |
| GONNET | 17 | W-C-G-P-H-N-D-E-R-QK-A-ST-F-Y-V-M-IL | W-C-G-P-H-N-D-E-R-K-A-T-F-Y-V-M-L |
| Genetic algorithm | 5 | AG-C-DEKNPQRST-FILMVWY-H | G-C-P-Y-H |
| Hierarchical clustering | 5 | PWDHI-GYCS-REL-NF-TVKMAQ | I-S-L-F-A |
| Hierarchical clustering | 6 | STAND-G-RQEK-HP-IVLMWYF-C | N-G-E-H-Y-C |
| Hierarchical clustering | 7 | C-DE-RK-STNQ-HP-VILMF-WYAG | C-D-R-Q-H-M-G |
| Hierarchical clustering | 8 | G-IVWFY-ALM-EQRK-P-ND-C-HST | G-F-L-E-P-N-C-S |
| Hierarchical clustering | 9 | PW-DHI-GY-CS-REL-NF-TV-KM-AQ | P-H-G-S-E-F-V-M-Q |
| Hierarchical clustering | 10 | ST-A-ND-G-RQ-EK-HP-IVLM-WYF-C | T-A-D-G-Q-E-H-I-Y-C |
| Hierarchical clustering | 11 | RK-QE-ND-H-ST-P-A-G-IVLM-FYW-C | K-Q-D-H-T-P-A-G-V-F-C |
| Hierarchical clustering | 12 | ST-A-N-D-G-RQ-EK-H-P-IVLM-WYF-C | S-A-N-D-G-R-E-H-P-M-F-C |
| Hierarchical clustering | 13 | RK-QE-ND-H-S-T-P-A-G-IV-LM-FYW-C | R-E-D-H-S-T-P-A-G-V-M-Y-C |
| Hierarchical clustering | 14 | RK-QE-N-D-H-S-T-P-A-G-IV-LM-FYW-C | K-E-N-D-H-S-T-P-A-G-I-L-F-C |
| Hierarchical clustering | 15 | C-DE-R-K-ST-NQ-H-P-VIL-M-F-W-Y-A-G | C-E-R-K-S-Q-H-P-L-M-F-W-Y-A-G |
| Hierarchical clustering | 16 | C-DE-R-K-ST-NQ-H-P-V-IL-M-F-W-Y-A-G | C-D-R-K-S-Q-H-P-V-L-M-F-W-Y-A-G |
| Hierarchical clustering | 17 | RK-Q-E-N-D-H-S-T-P-A-G-IV-L-M-FY-W-C | K-Q-E-N-D-H-S-T-P-A-G-V-L-M-Y-W-C |
| Hierarchical clustering | 18 | S-T-A-N-D-G-R-Q-E-K-H-P-IV-L-M-WY-F-C | S-T-A-N-D-G-R-Q-E-K-H-P-I-L-M-Y-F-C |
| Hierarchical clustering | 19 | S-T-A-N-D-G-R-Q-E-K-H-P-I-V-L-M-WY-F-C | S-T-A-N-D-G-R-Q-E-K-H-P-I-V-L-M-W-F-C |
| Information theory | 8 | DKR-EA-GP-STNQ-H-C-WY-FMLIV | R-A-G-T-H-C-Y-L |
| JTT | 5 | AGPST-CFWY-DEN-HKQR-ILMV | G-Y-N-K-M |
| JTT | 6 | APST-CW-DEGN-FHY-ILMV-KQR | S-C-N-F-M-K |
| JTT | 7 | AGST-CW-DEN-FY-HP-ILMV-KQR | G-C-D-F-H-M-K |
| JTT | 8 | AST-CG-DEN-FY-HP-ILV-KQR-MW | S-C-E-F-H-V-R-M |
| JTT | 9 | AST-CW-DE-FY-GN-HQ-ILV-KR-MP | T-C-D-Y-G-Q-I-R-M |
| JTT | 10 | AST-CW-DE-FY-GN-HQ-IV-KR-LM-P | T-W-D-Y-N-Q-V-K-M-P |
| JTT | 11 | AST-C-DE-FY-GN-HQ-IV-KR-LM-P-W | A-C-D-F-N-H-I-R-L-P-W |
| JTT | 12 | AST-C-DE-FY-G-HQ-IV-KR-LM-N-P-W | A-C-D-Y-G-H-V-K-L-N-P-W |
| JTT | 13 | AST-C-DE-FY-G-H-IV-KR-LM-N-P-Q-W | S-C-E-Y-G-H-V-R-L-N-P-Q-W |
| JTT | 14 | AST-C-DE-FL-G-H-IV-KR-M-N-P-Q-W-Y | T-C-E-F-G-H-V-R-M-N-P-Q-W-Y |
| JTT | 15 | AST-C-DE-F-G-H-IV-KR-L-M-N-P-Q-W-Y | T-C-D-F-G-H-I-K-L-M-N-P-Q-W-Y |
| JTT | 16 | AT-C-DE-F-G-H-IV-KR-L-M-N-P-Q-S-W-Y | A-C-D-F-G-H-I-R-L-M-N-P-Q-S-W-Y |
| JTT | 17 | AT-C-DE-F-G-H-IV-K-L-M-N-P-Q-R-S-W-Y | A-C-E-F-G-H-V-K-L-M-N-P-Q-R-S-W-Y |
| JTT | 18 | A-C-DE-F-G-H-IV-K-L-M-N-P-Q-R-S-T-W-Y | A-C-E-F-G-H-I-K-L-M-N-P-Q-R-S-T-W-Y |
| JTT | 19 | A-C-D-E-F-G-H-IV-K-L-M-N-P-Q-R-S-T-W-Y | A-C-D-E-F-G-H-I-K-L-M-N-P-Q-R-S-T-W-Y |
| K-means | 8 | ED-QST-NH-YPRK-LMI-VAF-GW-C | D-T-H-R-M-V-W-C |
| MIG | 5 | G-DN-AEHKQRST-CFILMVWY-P | G-D-T-C-P |
| MIG | 6 | G-DN-AEFHILMKQRVWY-CT-S-P | G-N-H-T-S-P |

| | | | |
|---|---|---|---|
| MIG | 7 | G-DN-AEFILMKQRVWY-CH-T-S-P | G-N-E-C-T-S-P |
| MIG | 8 | G-D-N-AEFILMKQRVWY-CH-T-S-P | G-D-N-W-C-T-S-P |
| MIG | 9 | G-D-N-AEFILMKQRVWY-C-H-T-S-P | G-D-N-Q-C-H-T-S-P |
| MIG | 10 | G-D-N-AEFILMKQRVW-Y-C-H-T-S-P | G-D-N-F-Y-C-H-T-S-P |
| MIG | 11 | G-D-N-AEFILMKQRV-W-Y-C-H-T-S-P | G-D-N-V-W-Y-C-H-T-S-P |
| MIG | 12 | G-D-N-AEFILMKQV-R-W-Y-C-H-T-S-P | G-D-N-E-R-W-Y-C-H-T-S-P |
| MIG | 13 | G-D-N-AEFILMKV-Q-R-W-Y-C-H-T-S-P | G-D-N-K-Q-R-W-Y-C-H-T-S-P |
| MIG | 14 | G-D-N-AEFILKV-M-Q-R-W-Y-C-H-T-S-P | G-D-N-I-M-Q-R-W-Y-C-H-T-S-P |
| MJ | 5 | MFILV-AWC-YQHPGTSN-RK-DE | V-A-H-R-E |
| MJ | 6 | MFILV-A-C-WYQHPGTSN-RK-DE | I-A-C-G-K-E |
| MJ | 7 | MFILV-A-C-WYQHP-GTSN-RK-DE | F-A-C-Y-N-R-E |
| MJ | 8 | LF-I-MVW-CY-HAT-GP-RQS-NEDK | L-I-V-Y-H-P-S-D |
| MJ | 9 | MF-ILV-A-C-WYQHP-G-TSN-RK-DE | F-V-A-C-P-G-S-R-D |
| MJ | 10 | QSNTGA-P-ED-LIVM-FW-Y-H-R-K-C | G-P-D-I-W-Y-H-R-K-C |
| MJ | 11 | MF-IL-V-A-C-WYQHP-G-TSN-RK-D-E | F-I-V-A-C-W-G-N-R-D-E |
| MJ | 12 | MF-IL-V-A-C-WYQHP-G-TS-N-RK-D-E | F-L-V-A-C-P-G-T-N-K-D-E |
| MJ | 13 | QSNT-G-A-P-E-D-LIVM-FW-Y-H-R-K-C | N-G-A-P-E-D-L-W-Y-H-R-K-C |
| MJ | 14 | QSNT-G-A-P-E-D-LIVM-F-W-Y-H-R-K-C | N-G-A-P-E-D-M-F-W-Y-H-R-K-C |
| MJ | 15 | QSN-T-G-A-P-E-D-LIVM-F-W-Y-H-R-K-C | Q-T-G-A-P-E-D-M-F-W-Y-H-R-K-C |
| MJ | 16 | MF-I-L-V-A-C-WYQ-H-P-G-T-S-N-RK-D-E | F-I-L-V-A-C-Q-H-P-G-T-S-N-R-D-E |
| MJ | 18 | Q-S-N-T-G-A-P-E-D-LIV-F-M-W-Y-H-R-K-C | Q-S-N-T-G-A-P-E-D-V-F-M-W-Y-H-R-K-C |
| MJ | 19 | Q-S-N-T-G-A-P-E-D-LI-V-F-M-W-Y-H-R-K-C | Q-S-N-T-G-A-P-E-D-I-V-F-M-W-Y-H-R-K-C |
| MJ & BLOSUM62 | 13 | MF-IL-V-A-C-WYQHP-G-T-S-N-RK-D-E | F-I-V-A-C-P-G-T-S-N-K-D-E |
| MJ & BLOSUM62 | 14 | IMV-L-F-WY-G-P-C-A-S-T-N-HRKQ-E-D | V-L-F-W-G-P-C-A-S-T-N-K-E-D |
| MJ & BLOSUM62 | 19 | P-G-E-K-R-Q-D-S-N-T-H-C-I-V-W-YF-A-L-M | P-G-E-K-R-Q-D-S-N-T-H-C-I-V-W-Y-A-L-M |
| MMI | 5 | FWY-CILMV-DEGKNS-APQT-HR | Y-I-D-A-R |
| MMI | 6 | FWY-CILMV-DE-GKNQS-APT-HR | F-L-D-S-T-R |
| MMI | 7 | FWY-CILMV-DE-K-GNPQS-AT-HR | Y-V-D-K-G-A-H |
| MMI | 8 | FWY-ILMV-C-DE-K-GNPQS-AT-HR | W-L-C-D-K-P-T-R |
| MMI | 9 | FWY-ILMV-C-DE-K-GNQS-PT-A-HR | Y-M-C-E-K-S-T-A-H |
| MMI | 10 | WY-F-ILMV-C-DE-K-GNQS-PT-A-HR | W-F-V-C-D-K-N-T-A-R |
| MMI | 11 | WY-F-ILMV-C-DE-K-G-PNQS-T-A-HR | W-F-L-C-E-K-G-Q-T-A-R |
| MMI | 12 | WY-F-IL-MV-C-DE-K-G-PNQS-T-A-HR | Y-F-I-M-C-D-K-G-S-T-A-R |
| MMI | 13 | WY-F-IL-MV-C-DE-K-G-P-NQS-T-A-HR | Y-F-I-M-C-D-K-G-P-Q-T-A-H |
| MMI | 14 | W-Y-F-IL-MV-C-DE-K-G-P-NQS-T-A-HR | W-Y-F-L-V-C-E-K-G-P-S-T-A-H |
| MMI | 15 | W-Y-F-IL-MV-C-DE-K-G-P-NQS-T-A-H-R | W-Y-F-I-M-C-E-K-G-P-S-T-A-H-R |
| MMI | 16 | W-Y-F-IL-M-V-C-DE-K-G-P-NQS-T-A-H-R | W-Y-F-I-M-V-C-D-K-G-P-S-T-A-H-R |
| MMI | 17 | W-Y-F-I-L-M-V-C-DE-K-G-P-NQS-T-A-H-R | W-Y-F-I-L-M-V-C-D-K-G-P-N-T-A-H-R |
| MMI | 18 | W-Y-F-I-L-M-V-C-DE-K-G-P-N-QS-T-A-H-R | W-Y-F-I-L-M-V-C-D-K-G-P-N-S-T-A-H-R |
| MMI | 19 | W-Y-F-I-L-M-V-C-D-E-K-G-P-N-QS-T-A-H-R | W-Y-F-I-L-M-V-C-D-E-K-G-P-N-Q-T-A-H-R |
| PAM | 5 | AGSPDEQNHTKR-MIVL-FY-C-W | H-V-Y-C-W |
| PAM | 6 | AGSP-DEQNHTKR-MIL-FY-CV-W | S-K-L-Y-V-W |
| PAM | 7 | AGP-DEQNH-TKRIMV-L-FY-CS-W | G-D-M-L-F-C-W |

| | | | |
|---|---|---|---|
| PAM | 8 | AG-DEQN-TKRIMV-L-HY-CS-FP-W | A-E-V-L-H-S-P-W |
| PAM | 9 | AG-P-DEQN-TKRIM-L-F-HY-VCS-W | G-P-E-K-L-F-H-S-W |
| PAM | 10 | AG-P-DEQN-TKRM-L-F-HY-VCS-I-W | G-P-Q-K-L-F-H-C-I-W |
| PAM | 11 | AG-P-DEQN-TK-RI-ML-F-H-Y-VCS-W | G-P-Q-T-R-L-F-H-Y-S-W |
| PAM | 12 | FAS-P-G-DEQ-NL-TK-R-H-W-Y-IM-VC | F-P-G-D-L-T-R-H-W-Y-M-V |
| PAM | 13 | FAS-P-G-DEQ-NL-T-K-R-H-W-Y-IM-VC | S-P-G-E-N-T-K-R-H-W-Y-M-V |
| PAM | 14 | FA-P-G-T-DE-QM-NL-K-R-H-W-Y-IV-SC | F-P-G-T-D-M-N-K-R-H-W-Y-I-S |
| PAM | 15 | FAS-P-G-DE-T-Q-NL-K-R-H-W-Y-I-M-VC | S-P-G-D-T-Q-N-K-R-H-W-Y-I-M-C |
| PAM | 16 | FA-P-G-ST-DE-Q-N-K-R-H-W-Y-M-L-I-VC | A-P-G-S-E-Q-N-K-R-H-W-Y-M-L-I-V |
| PAM | 17 | FA-P-G-ST-DE-Q-N-K-R-H-W-Y-M-L-I-V-C | F-P-G-T-D-Q-N-K-R-H-W-Y-M-L-I-V-C |
| PAM | 18 | FA-P-G-S-T-DE-Q-N-K-R-H-W-Y-M-L-I-V-C | F-P-G-S-T-D-Q-N-K-R-H-W-Y-M-L-I-V-C |
| PAM | 19 | FA-P-G-S-T-D-E-Q-N-K-R-H-W-Y-M-L-I-V-C | F-P-G-S-T-D-E-Q-N-K-R-H-W-Y-M-L-I-V-C |
| PSO | 5 | EL-DTV-RGKFW-AQHIPY-NCMS | L-T-R-P-N |
| PSO | 6 | CILKMSV-Y-FW-G-R-ANDQEHPT | L-Y-F-G-R-T |
| PSO | 8 | NQEHKPS-L-WY-R-I-G-V-ADCMFT | S-L-W-R-I-G-V-C |
| Physical-chemical | 6 | RDENQKH-LIVAMF-STYW-P-G-C | E-I-Y-P-G-C |
| Physical-chemical | 7 | AV-CGNP-D-EKRQ-FWYH-ILM-ST | V-P-D-E-F-M-T |
| Physical-chemical | 9 | AV-CGNP-D-EKR-Q-FWY-H-ILM-ST | A-C-D-K-Q-Y-H-L-T |
| Physical-chemical | 11 | AV-C-GNP-D-EKR-Q-FWY-H-IL-M-ST | A-C-G-D-E-Q-F-H-I-M-S |
| Physical-chemical | 19 | G-I-LV-F-Y-W-A-M-E-Q-R-K-P-N-D-H-S-T-C | G-I-V-F-Y-W-A-M-E-Q-R-K-P-N-D-H-S-T-C |
| Protein blocks | 5 | G-IVFYW-ALMEQRK-P-NDHSTC | G-V-M-P-N |
| Protein blocks | 8 | DKR-EA-GP-STNQ-H-C-WY-FMLIV | R-E-G-T-H-C-Y-L |
| Protein blocks | 9 | G-IV-FYW-ALM-EQRK-P-ND-HS-TC | G-I-F-M-E-P-D-S-T |
| Protein blocks | 10 | DNS-EKRQ-TH-GP-AM-C-W-F-YL-IV | N-E-T-G-A-C-W-F-L-I |
| Protein blocks | 11 | G-IV-FYW-A-LM-EQRK-P-ND-HS-T-C | G-I-F-A-L-R-P-N-S-T-C |
| Protein blocks | 13 | G-IV-FYW-A-L-M-E-QRK-P-ND-HS-T-C | G-V-W-A-L-M-E-K-P-N-H-T-C |
| SDM | 5 | P-HGAREKQSTND-W-C-MVILFY | P-H-W-C-F |
| SDM | 6 | P-H-GAREKQSTND-W-C-MVILFY | P-H-E-W-C-I |
| SDM | 7 | P-H-G-AREKQSTND-W-C-MVILFY | P-H-G-D-W-C-V |
| SDM | 8 | P-H-G-AREKQST-ND-W-C-MVILFY | P-H-G-E-D-W-C-F |
| SDM | 9 | P-H-G-AREKQST-ND-W-C-MVIL-FY | P-H-G-A-N-W-C-V-F |
| SDM | 10 | P-H-G-A-REKQST-ND-W-C-MVIL-FY | P-H-G-A-S-D-W-C-M-F |
| SDM | 11 | P-H-G-A-REK-QST-ND-W-C-MVIL-FY | P-H-G-A-R-T-D-W-C-I-F |
| SDM | 12 | P-H-G-A-REK-QST-N-D-W-C-MVIL-FY | P-H-G-A-E-Q-N-D-W-C-I-F |
| SDM | 13 | P-H-G-A-REK-QST-N-D-W-C-MVIL-F-Y | P-H-G-A-K-Q-N-D-W-C-I-F-Y |
| SDM | 14 | P-H-G-A-REK-QST-N-D-W-C-M-VIL-F-Y | P-H-G-A-R-T-N-D-W-C-M-V-F-Y |
| SDM | 15 | P-H-G-A-R-EK-QST-N-D-W-C-M-VIL-F-Y | P-H-G-A-R-K-Q-N-D-W-C-M-V-F-Y |
| SDM | 16 | P-H-G-A-R-EK-Q-ST-N-P-W-C-M-VIL-F-Y | P-H-G-A-R-E-Q-S-N-P-W-C-M-V-F-Y |
| SDM | 17 | P-H-G-A-R-EK-Q-S-T-N-P-W-C-M-VIL-F-Y | P-H-G-A-R-E-Q-S-T-N-P-W-C-M-I-F-Y |
| Sequence alignments | 5 | DSHFM-ERQL-KPAC-NTWY-GIV | H-R-K-T-G |
| Sequence alignments | 6 | DENQ-KRH-STGPA-C-WYF-MLIV | N-R-T-C-W-M |
| Sequence alignments | 10 | DN-EQ-KR-STA-G-P-HW-C-YF-MLIV | N-Q-K-S-G-P-W-C-Y-L |
| Sequence alignments | 13 | D-E-KQR-NS-T-G-P-H-A-C-WYF-ML-IV | D-E-Q-N-T-G-P-H-A-C-Y-M-V |

| | | | |
|---|---|---|---|
| Sequence alignments | 16 | D-E-N-KR-Q-ST-G-P-H-A-C-W-Y-F-ML-IV | D-E-N-K-Q-T-G-P-H-A-C-W-Y-F-M-V |
| Structure | 5 | PG-EKRQ-DSNTHC-IVWYF-ALM | P-K-S-I-A |
| Structure | 6 | PG-EKRQ-DSN-THC-IVWYF-ALM | P-Q-N-C-Y-L |
| Structure | 7 | PG-EKRQ-DSN-THC-IVWYF-A-LM | P-Q-D-C-I-A-M |
| Structure | 8 | P-G-EKRQ-DSN-THC-IVWYF-A-LM | P-G-R-N-C-V-A-M |
| Structure | 9 | P-G-EKRQ-DSN-THC-IV-WYF-A-LM | P-G-K-N-C-I-W-A-M |
| Structure | 10 | P-G-EKRQ-DSN-T-HC-IV-WYF-A-LM | P-G-R-D-T-H-I-W-A-L |
| Structure | 11 | P-G-EKRQ-D-SN-T-HC-IV-WYF-A-LM | P-G-Q-D-N-T-C-I-F-A-M |
| Structure | 12 | P-G-EKRQ-D-SN-T-H-C-IV-WYF-A-LM | P-G-K-D-N-T-H-C-I-W-A-L |
| Structure | 13 | P-G-E-KRQ-D-SN-T-H-C-IV-WYF-A-LM | P-G-E-K-D-N-T-H-C-I-F-A-L |
| Structure | 15 | P-G-E-KRQ-D-S-N-T-H-C-IV-W-YF-A-LM | P-G-E-R-D-S-N-T-H-C-V-W-F-A-M |
| Structure | 16 | P-G-E-KRQ-D-S-N-T-H-C-IV-W-YF-A-L-M | P-G-E-Q-D-S-N-T-H-C-I-W-F-A-L-M |
| Structure | 17 | P-G-E-K-RQ-D-S-N-T-H-C-IV-W-YF-A-L-M | P-G-E-K-R-D-S-N-T-H-C-V-W-F-A-L-M |
| Structure | 18 | P-G-E-K-RQ-D-S-N-T-H-C-I-V-W-YF-A-L-M | P-G-E-K-R-D-S-N-T-H-C-I-V-W-Y-A-L-M |
| Structure | 19 | P-G-E-K-R-Q-D-S-N-T-H-C-I-V-W-YF-A-L-M | P-G-E-K-R-Q-D-S-N-T-H-C-I-V-W-F-A-L-M |
| Structure alignments | 5 | DE-NKRQS-THA-GP-CWYFMLIV | E-R-H-G-V |
| Structure alignments | 6 | DE-KR-NQST-GP-HWYF-ACMLIV | D-R-N-G-H-M |
| Structure alignments | 12 | D-N-EKR-QST-G-P-H-A-C-W-YF-MLIV | D-N-E-S-G-P-H-A-C-W-F-L |
| Structure alignments | 15 | D-N-E-KRQ-S-T-G-P-H-A-C-W-YF-ML-IV | D-N-E-R-S-T-G-P-H-A-C-W-Y-M-I |
| Structure alignments | 17 | D-EK-N-R-Q-S-T-G-P-H-A-C-W-Y-F-M-LIV | D-K-N-R-Q-S-T-G-P-H-A-C-W-Y-F-M-L |
| UPGMA | 5 | ASTNDRQEKIVLMGP-FY-H-C-W | S-Y-H-C-W |
| UPGMA | 6 | ASTNDRQEKIVLMG-P-FY-H-C-W | S-P-F-H-C-W |
| UPGMA | 7 | ASTNDRQEKIVLM-G-P-FY-H-C-W | N-G-P-F-H-C-W |
| UPGMA | 8 | ASTNDRQEK-IVLM-G-P-FY-H-C-W | E-V-G-P-Y-H-C-W |
| UPGMA | 10 | ASTND-RQEK-IVLM-G-P-F-Y-H-C-W | T-Q-I-G-P-F-Y-H-C-W |
| UPGMA | 11 | ASTN-D-RQEK-IVLM-G-P-F-Y-H-C-W | S-D-Q-V-G-P-F-Y-H-C-W |
| UPGMA | 12 | ASTN-D-RQ-EK-IVLM-G-P-F-Y-H-C-W | A-D-R-E-M-G-P-F-Y-H-C-W |
| UPGMA | 14 | AST-N-D-RQ-EK-IVL-M-G-P-F-Y-H-C-W | T-N-D-R-K-V-M-G-P-F-Y-H-C-W |
| UPGMA | 15 | AST-N-D-R-Q-EK-IVL-M-G-P-F-Y-H-C-W | S-N-D-R-Q-E-V-M-G-P-F-Y-H-C-W |
| UPGMA | 16 | AST-N-D-R-Q-E-K-IVL-M-G-P-F-Y-H-C-W | T-N-D-R-Q-E-K-L-M-G-P-F-Y-H-C-W |
| UPGMA | 17 | AST-N-D-R-Q-E-K-IV-L-M-G-P-F-Y-H-C-W | S-N-D-R-Q-E-K-I-L-M-G-P-F-Y-H-C-W |
| UPGMA | 18 | AS-T-N-D-R-Q-E-K-IV-L-M-G-P-F-Y-H-C-W | S-T-N-D-R-Q-E-K-I-L-M-G-P-F-Y-H-C-W |
| UVG | 5 | LIVGAP-QNMTSC-ED-KR-YFWH | A-Q-E-R-W |
| UVG | 6 | LIVGAP-QNMTSC-ED-KR-YFW-H | P-S-E-K-W-H |
| UVG | 7 | LIVGAP-QNM-TSC-ED-KR-YFW-H | A-N-T-E-K-W-H |
| UVG | 8 | LIV-GAP-QNM-TSC-ED-KR-YFW-H | V-P-M-T-E-R-Y-H |
| UVG | 10 | LIV-GA-P-QN-M-TSC-ED-KR-YFW-H | I-A-P-Q-M-S-D-K-W-H |
| UVG | 13 | LIV-GA-P-QN-M-TS-C-ED-KR-Y-F-W-H | I-A-P-Q-M-T-C-E-K-Y-F-W-H |
| UVG | 14 | LIV-GA-P-QN-M-TS-C-ED-K-R-Y-F-W-H | I-A-P-N-M-S-C-D-K-R-Y-F-W-H |
| UVG | 15 | LIV-GA-P-QN-M-T-S-C-ED-K-R-Y-F-W-H | I-A-P-N-M-T-S-C-E-K-R-Y-F-W-H |
| UVG | 16 | LIV-GA-P-Q-N-M-T-S-C-ED-K-R-Y-F-W-H | L-A-P-Q-N-M-T-S-C-D-K-R-Y-F-W-H |
| UVG | 17 | LIV-G-A-P-Q-N-M-T-S-C-ED-K-R-Y-F-W-H | I-G-A-P-Q-N-M-T-S-C-E-K-R-Y-F-W-H |
| UVG | 18 | LI-V-G-A-P-Q-N-M-T-S-C-ED-K-R-Y-F-W-H | I-V-G-A-P-Q-N-M-T-S-C-E-K-R-Y-F-W-H |

| | | | |
|---|---|---|---|
| UVG | 19 | LI-V-G-A-P-Q-N-M-T-S-C-E-D-K-R-Y-F-W-H | L-V-G-A-P-Q-N-M-T-S-C-E-D-K-R-Y-F-W-H |
| Unweighted pair group | 5 | AWMGST-HPY-CVIFL-DNQ-ERK | T-H-C-D-R |
| Unweighted pair group | 6 | AGTLV-DKNFQ-EIPRS-CMY-H-W | V-Q-E-M-H-W |
| Unweighted pair group | 7 | AEM-L-CDT-NY-FIKHQVRW-GS-P | M-L-D-Y-H-G-P |
| Unweighted pair group | 8 | AWM-GST-HPY-CVI-FL-DNQ-ER-K | A-T-Y-I-L-Q-R-K |
| Unweighted pair group | 9 | AWM-GS-T-HPY-CVI-FL-DNQ-ER-K | A-G-T-Y-C-L-N-R-K |
| Unweighted pair group | 10 | AEM-L-CDT-NY-FIKH-QV-RW-G-S-P | M-L-D-Y-K-Q-W-G-S-P |
| Unweighted pair group | 11 | A-GT-LV-DKN-FQ-E-IPR-S-CMY-H-W | A-T-L-D-F-E-R-S-C-H-W |
| Unweighted pair group | 12 | AWM-G-S-T-H-PY-CVI-FL-DNQ-E-R-K | M-G-S-T-H-Y-V-F-Q-E-R-K |
| Unweighted pair group | 13 | A-G-T-LV-DKN-FQ-E-IPR-S-CM-Y-H-W | A-G-T-L-N-Q-E-P-S-M-Y-H-W |
| Unweighted pair group | 14 | A-EM-L-C-DT-NY-FIK-H-Q-V-RW-G-S-P | A-E-L-C-D-Y-F-H-Q-V-W-G-S-P |
| Unweighted pair group | 15 | A-C-V-HP-L-D-Q-S-ER-GN-F-IMT-K-W-Y | A-C-V-H-L-D-Q-S-E-G-F-I-K-W-Y |
| Unweighted pair group | 16 | A-C-V-HP-L-D-Q-S-ER-GN-F-I-MT-K-W-Y | A-C-V-P-L-D-Q-S-E-N-F-I-M-K-W-Y |
| Unweighted pair group | 17 | A-C-V-H-P-L-D-Q-S-ER-GN-F-I-MT-K-W-Y | A-C-V-H-P-L-D-Q-S-R-G-F-I-T-K-W-Y |
| Unweighted pair group | 18 | A-G-T-L-V-DK-N-F-Q-E-I-P-R-S-CM-Y-H-W | A-G-T-L-V-D-N-F-Q-E-I-P-R-S-C-Y-H-W |
| Unweighted pair group | 19 | A-E-M-L-C-D-T-NY-F-I-K-H-Q-V-R-W-G-S-P | A-E-M-L-C-D-T-N-F-I-K-H-Q-V-R-W-G-S-P |
| WAG | 5 | HRKQNEDSTGPA-CV-IML-FY-W | E-V-M-Y-W |
| WAG | 6 | HRKQNEDSTPA-G-CV-IML-FY-W | H-G-V-I-Y-W |
| WAG | 7 | HRKQNEDSTA-G-P-CV-IML-FY-W | K-G-P-V-I-F-W |
| WAG | 8 | HRKQSTA-NED-G-P-CV-IML-FY-W | K-E-G-P-C-L-Y-W |
| WAG | 9 | HRKQ-NED-ASTG-P-C-IV-MLF-Y-W | K-E-G-P-C-V-M-Y-W |
| WAG | 10 | HRKSA-Q-NED-G-P-C-TIV-MLF-Y-W | H-Q-D-G-P-C-I-F-Y-W |
| WAG | 11 | RKQ-NG-ED-AST-P-C-IV-HML-F-Y-W | R-G-E-A-P-C-I-H-F-Y-W |
| WAG | 12 | RKQ-ED-NAST-G-P-C-IV-H-ML-F-Y-W | K-D-S-G-P-C-I-H-L-F-Y-W |
| WAG | 13 | RK-QE-D-NG-HA-ST-P-C-IV-ML-F-Y-W | K-Q-D-G-H-S-P-C-V-M-F-Y-W |
| WAG | 14 | R-K-QE-D-NG-HA-ST-P-C-IV-ML-F-Y-W | R-K-Q-D-G-H-S-P-C-V-L-F-Y-W |
| WAG | 15 | R-K-QE-D-NG-HA-ST-P-C-IV-M-L-F-Y-W | R-K-Q-D-G-H-T-P-C-I-M-L-F-Y-W |
| WAG | 16 | R-K-Q-E-D-NG-HA-ST-P-C-IV-M-L-F-Y-W | R-K-Q-E-D-G-H-T-P-C-I-M-L-F-Y-W |
| WAG | 17 | R-K-Q-E-D-NG-HA-S-T-P-C-IV-M-L-F-Y-W | R-K-Q-E-D-N-H-S-T-P-C-I-M-L-F-Y-W |
| WAG | 18 | R-K-Q-E-D-NG-HA-S-T-P-C-I-V-M-L-F-Y-W | R-K-Q-E-D-G-H-S-T-P-C-I-V-M-L-F-Y-W |
| WAG | 19 | R-K-Q-E-D-NG-H-A-S-T-P-C-I-V-M-L-F-Y-W | R-K-Q-E-D-G-H-A-S-T-P-C-I-V-M-L-F-Y-W |
| variance maximization | 5 | WFYH-MLIV-GP-CATS-NDEQRK | W-V-P-A-K |
| variance maximization | 6 | WFYH-MLIV-G-P-CATS-NDEQRK | W-L-G-P-A-R |
| variance maximization | 7 | WFY-MLIV-G-P-CATS-NDE-HQRK | W-V-G-P-S-E-H |
| variance maximization | 8 | WFY-MLIV-C-G-P-ATS-NDE-HQRK | W-L-C-G-P-S-D-H |
| variance maximization | 9 | WFY-MLIV-C-G-P-ATS-NDE-H-QRK | Y-L-C-G-P-T-E-H-K |
| variance maximization | 10 | W-FY-MLIV-C-G-P-ATS-NDE-H-QRK | W-F-M-C-G-P-T-E-H-R |
| variance maximization | 11 | W-FY-MLIV-C-G-P-ATS-N-DE-H-QRK | W-F-I-C-G-P-T-N-D-H-Q |
| variance maximization | 12 | W-FY-MLIV-A-C-G-P-TS-Q-NDE-H-RK | W-Y-L-A-C-G-P-T-Q-E-H-R |
| variance maximization | 13 | WFY-MLIV-A-C-G-P-TS-N-Q-D-E-H-RK | Y-L-A-C-G-P-S-N-Q-D-E-H-K |
| variance maximization | 14 | W-FY-MLIV-A-C-G-P-TS-N-Q-D-E-H-RK | W-F-M-A-C-G-P-T-N-Q-D-E-H-R |
| variance maximization | 15 | W-FY-M-LIV-A-C-G-P-TS-N-Q-D-E-H-RK | W-Y-M-I-A-C-G-P-T-N-Q-D-E-H-K |
| variance maximization | 16 | W-F-Y-M-LIV-A-C-G-P-TS-N-Q-D-E-H-RK | W-F-Y-M-L-A-C-G-P-S-N-Q-D-E-H-R |

| | | | |
|---|---|---|---|
| variance maximization | 17 | W-F-Y-M-LIV-A-C-G-P-T-S-N-Q-D-E-H-RK | W-F-Y-M-V-A-C-G-P-T-S-N-Q-D-E-H-K |
| variance maximization | 18 | W-F-Y-M-LIV-A-C-G-P-T-S-N-Q-D-E-H-R-K | W-F-Y-M-L-A-C-G-P-T-S-N-Q-D-E-H-R-K |
| variance maximization | 19 | W-F-Y-M-L-IV-A-C-G-P-T-S-N-Q-D-E-H-R-K | W-F-Y-M-L-I-A-C-G-P-T-S-N-Q-D-E-H-R-K |
